# Competition, cooperation and immune selection of multi-strain Plasmodium falciparum malaria

**DOI:** 10.1101/539676

**Authors:** David Gurarie

## Abstract

**Setup:** Malaria Plasmodium falciparum (Pf) species contains multiple strains with different immunogenic profiles, and expressed phenotypes. These strains circulate in host populations via mosquito transmission, and interact (compete, cooperate) on two levels: within - host (via cross-reactive immunity), and in host populations. Both factors, host immunity and transmission environment, play important part in evolution and selection.

Conventional population-based models of malaria have limited capacity to accommodate parasite-immune dynamics within-host and strain diversity. Here we developed an in-host model for multi-strain malaria based on its genetic (immunogenic) makeup, which accounts for essential parasite-immune biology. The model allows efficient simulations of mixed-strain infections in individual hosts and in host ensembles over multiple transmission cycles. We use it to explore evolutionary implications (competition, selection) of malaria quasi-species, driven by host immunity and transmission intensity.

**Results:** The key ‘selectable’ trait within-host is strain *transmissibility* (TP), which measures cumulative odds of mosquito infection by a given strain over infection history. Here we adopt it to explore evolutionary implications of parasite-immune interactions on different time scales and transmission environments. Specifically, we explore (i) *primary strain selection* in naïve host ensembles based on TP-fitness; (ii) *evolution* and *selection* of mixed multi-strain systems over *multiple transmission cycles*.

On level (i) different strain mixtures competed in multiple hosts, to identify ‘most fit’ (highly transmissible) types. A key observation of (i) was *fitness-cost* of in-host competition, i.e. statistical TP-loss determined by multiplicity of infection (number of competing strains), and strain genotype (immunogenic profile). The most-fit strains maintained their high TP-values regardless of competing environment.

We selected them for step (ii), to explore long-term evolution over multiple transmission cycles. Our analysis revealed peculiar features of evolution: success within-host (step (i)) did not guarantee strain survival over multiple cycles. Indeed, the latter was strongly associated with *cooperative behavior*, i.e. co-existence of a given strain in suitable mixtures, in multiple hosts over many generations. We examined the resulting population structure of evolving strains, in terms of their immune cross-reactivity. Overall, our results were consistent with predictions of *strain theory* [1–4], [5, 6]. Strain theory predicts that cross-reacting parasite strains in host population should organize themselves into ‘non-overlapping’ (immunogenically disjunct) clusters. In our case, no strict ‘immune separation’ arises, but cross-reactivity is lost over multiple cycles, and surviving clusters are ‘nearly disjunct’. Such weakly overlapping clusters (*cooperating cliques*) persisted over long (evolutionary) periods. Specifically, each clique was found to possess a *core node* -highly cooperative *persistent strain*, carrying a subordinate (transient) cluster.

Our results shed new light on relative importance of *competitive* vs. *cooperative* behavior, and multi-level organization of genetically structured parasite system. They could have implications for malaria control and vaccine design.

## Introduction

Multiple malaria strains circulate in host populations. Their makeup and population structure have important implications for parasite persistence, spread and control outcomes.

A successful parasite must succeed on two levels: (i) within-host, subject to immune pressure; (ii) in host communities and transmission environment (e.g. in vector stage). In case of malaria, mosquito vector – the definitive host, also serves to diversify parasite pool via mating and recombination. Strain diversity can contribute to parasite survival and adaption, particularly under stressful conditions.

There are different ways to define and quantify *strain diversity* (see e.g. [7]). Typically, it refers to internal stratification of species into distinct subgroups (types), based on their *phenotypes* or the underlying *genotype*. Many modeling approaches to host-parasite evolution (e.g. [8–11], [12], [13–15], [16]) employ hypothetical ‘fitness traits’, like virulence, transmissibility, persistence, and classify strains accordingly (‘virulent’ - ‘benign’, ‘drug sensitive’ - ‘tolerant’, et al). The underlying genetic makeup may not enter such models explicitly. Indeed, links between *genetic makeup* and *phenotypic expression* are often tenuous, confounded by multiple factors, including host immunity (status, competence), parasite strain mixture, timing of inoculae et al.

While host immunity is recognized as an important factor of parasite evolution, combining it with population-level transmission raises many modeling challenges, particularly for genetically diverse parasite populations and dynamic immune environment. Conventional population-based modeling approaches typically oversimplifies intrahost immunology, parasite (strain) makeup and immune regulation mechanisms (see e.g. [17],[18] [8–11], [12], [13–15], [16]). Their scope is limited, particularly for highly dynamic infections, and genetically diverse parasites, like malaria.

Individual-based modeling (IBM) offers greater flexibility and potential to resolve details of in-host biology and parasite-immune interactions (see e.g. [19–23], [16, 24, 25]). The challenge is to make such systems computationally efficient and scalable to population level transmission. Here we develop an IBM approach for multi-strain malaria, patterned after Pf parasite, which contains sufficient biological details, yet is computationally efficient to allow large host ensembles (communities) long infection histories, and multiple transmission cycles.

Our setup highlights genetically structured parasite systems, their blood stages (merozoite, gametocyte), invasion and depletion of red blood cells (RBC), innate and adaptive immunity, mosquito uptake of gametocytes, and possible recombination of parasite clones within mosquito.

We focus on asexual (haploid) parasite stages in human host, and define a Pf *strain* by its genetic makeup, specifically a combination of var-genes (see e.g. [20, 26, 27]). So each Pf clone consists of a collection of var-genes drawn from a hypothetical ‘natural pool’. Each var-gene is assigned its specific growth/proliferation factor, and immunogenic profile (stimulated antibodies), see [26, 27]. During asexual replication-cycle, parasites (merozoite stage) undergo antigenic variation by expressing different var-genes on the surface of infected RBC. Mixed strains interact within-host through shared resource (RBC), and via innate and specific (cross-reactive) immunity. Transmission by random mosquito biting allows parasite recombination, production and release of new clones.

Evolution of host-parasite systems is often conceptualized via hypothetical *fitness traits*, e.g. virulence-transmissibility-persistence (V-T-P), whose trade-offs are believed to drive selection process (see e.g. [28–31]). In our setup, such fitness traits are not primary, but arise from complex parasite-immune interactions over the course of infection history. The resulting outcomes, strain-specific *virulence* (RBC depletion), *persistence* (detectable parasitemia), and *transmissibility* to mosquito, are highly variable depending on specifics of in-host (immune) environment, inoculation and infection dynamics.

The essential selectable trait on host-population level is strain *transmission potential* (TP) – cumulative probability of mosquito infection by a given strain over the course of infection. We use it extensively to identify most viable (transmissible) types in mixed infections, for naïve hosts (young children). Such TP-selection (we call it primary), gives a collection of strains surviving in young children, hence more likely to be observed and transmitted in older adults. We use them to study long-term parasite evolution in multiple transmission cycles (parasite generations).

For the long-term evolution study, we take most viable TP-types (highest quantile of the primary selection), and subject them (mixed-strain pools) to multiple transmission cycles via serial passage over naïve-hoist lines. We examine such evolving multi-strain systems, their var-gene makeup, cross-reactive immunogenic profiles (network structure), and population distribution in host communities.

Our analysis reveals some peculiar features of immune selection for malaria quasi-species. On individual-host level, most-viable clones are able to maintain their high TP-values against different competitors (mixed infections). However, not all such ‘competitive’ (high-TP) clones can succeed over multiple transmission cycles (host generation), and long evolutionary history. The long-term success required a pattern of cooperative behavior, whereby surviving clones would organize themselves into a cluster of ‘cooperating cliques’, with low immune cross-reactivity.

Strain diversity and population structure of Pf malaria was subject of several recent studies. An experimental approach based on clinical isolates [6], exhibits *serum dominance* networks with peculiar organization highlighting immune control. However, there was no clear identification of hypothetical ‘clones’ in those studies. A series of recent papers [2, 5, 32] undertook detailed molecular analysis of var-gene isolates collected from endemic areas. They revealed high diversity of Pf clones, and specifics of their organization in host population. The follow-up modeling analysis [1] confirmed highly non-random ‘clustered’ structure of multi-strain malaria systems, that bear qualitative resemblance to theoretical predictions of ‘strain theory’ ([13–15, 33, 34]).

Our paper contributes to this line of work. We add new perspectives on relative impact of within-host (immune-dominated) dynamics vs. mosquito transmission. We also explore the role of ‘competition’ vs. ‘cooperation’ among mixed clones, and multiple levels of selection, from individual var-genes, to viable clones, to ‘cooperating cliques’

The current analysis is confined to naive hosts and relatively short infection intra-host histories (based on initial inoculum). Mosquito recombination does not enter our analysis explicitly, though we allow random strain inputs into host communities, drawn from ‘natural reservoir’.

The future work will extend the scope of the current study to more realistic systems, with life-long infection histories, demographically and geographically structured host populations, and realistic mosquito transmission (c.f. [35]).

## Materials and methods

### Background

The key features of Pf asexual (merozoite) stage in human host are its multi-gene makeup, and immune evasion strategy via *antigenic variation* (AV) (see [34], [11, 24, 33, 36]). Each Pf parasite (haploid intra-host stage) carries multi-gene families of antigens (VSA). One of them EMPf1 (erythrocyte membrane protein 1), encoded by so-called var-genes, is expressed on the surface of infected red blood cells (iRBC). These antigens play double role. On the one hand, they expose parasite to adaptive host immunity by stimulating specific antibodies. On the other hand, they help parasite avoid spleen clearing by attaching to small capillary tissues (cytoadherence), which causes some forms of severe Pf pathogenesis, like cerebral and placental malaria. Each Pf parasite carries 50-60 var genes, but only one of them is expressed at any given time on the surface of the infected erythrocyte (iRBC). At the end of 48-hour replication cycle a new generation of merozoites released from the raptured iRBC, can switch their var-gene expression in the next generation of iRBC, thereby evading immune clearing by accumulated antibodies. The net effect of such AV switching is multiple waves of parasitemia, whereby a Pf-strain can extend its infection duration and increase transmission potential.

There are different ways to accommodate strain diversity and AV in within-host malaria models (see e.g. [11, 24, 33, 37]). Our approach follows recent papers ([26], [1]), but many details of the model setup and implementation differ.

### In-host model

Our model highlights multi-variant parasite makeup, based on var-genes. Each strain consists of several loci (1,2,…,m), filled with copies of var-gene alleles, drawn from a large (natural) variant pool *N* ≫ *m* (see Figure 1). Each var-gene has associated growth/replication factor r (= mean number of viable merozoites released by a schizont expressing this variant), and a specific antibody *u*_*r*_, stimulated by all strains (iRBC) which express variant *r*. Each parasite strain *s* = (*r*_1_, *r*_2_,…, *r*_*m*_) is identified with its var-gene makeup (c.f. [26]). The order of variants (*r*_*i*_) in the strain is also important, as we assume a sequential AV-pattern, starting with locus 1 (variant *r*_1_), switching to its neighboring loci to the right and left (*i* → *i* ±1). We also assumed all variants immunogenically distinct.

**Figure 1:**
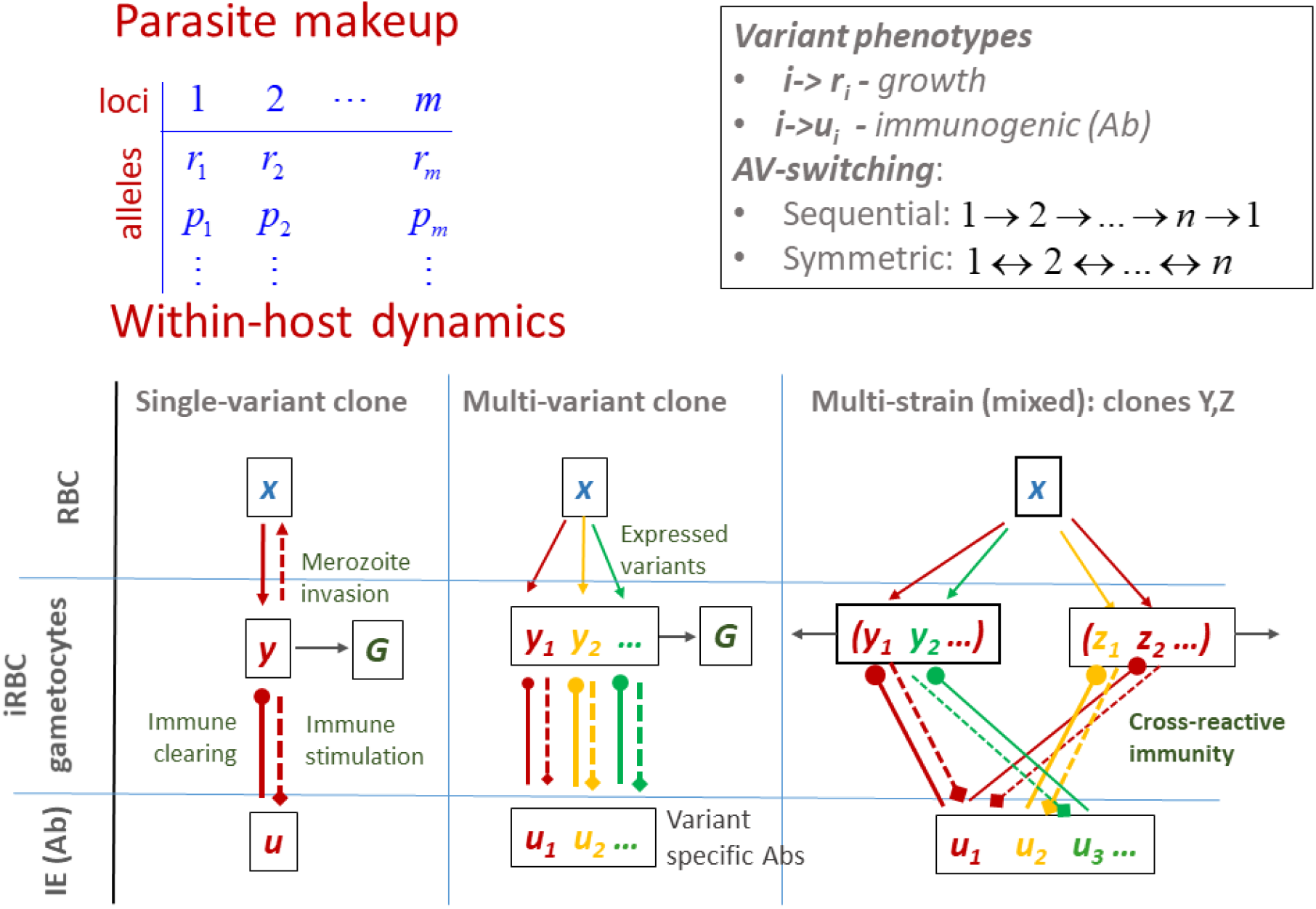
Schematic view of in-host model. Each strain is a combination of var-genes with specific growth and immunogenic profiles (antibodies), selected from large var-gene pool. The model features target cells (RBC), infected iRBC, that express different variants of a given strain, and stimulated specific immune effectors (IE). Shared variants (marked in different colors) create cross-reactivity patterns in mixed-strain infections. Different pattern of antigenic switching (AV) can be implemented in such setup.

So each parasite strain is characterized by its *immunogenic* and *growth* profiles, i.e. a suite of specific antibodies **u** = (*u*_1_,…, *u*_*m*_), and replication factors **r** = (*r*_1_,…, *r*_*m*_). Different clones can cross-react via shared var-genes, and stimulated Abs.

**Model** variables consist of cell populations (per muL of blood), and specific immune effectors identified with var-genes. Specifically we consider: (i) target RBC *x*, (ii) parasite populations (iRBC) 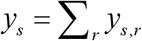, that carry strain *s*, each *y*_*s*_ is partitioned into subpopulations (*y*_*s*,*r*_) expressing variant *r*; (iii) gametocytemia *G*_*s*_ of strain s, (iv) immune effector variables {*u*_*r*_}, labeled by variants {*r*}.

In numeric simulations and plots (below) all cell populations (*x*, *y*, *G*) were rescaled relative to the normal RBC count, *x*_0_ = 5 ⋅10^6^ / *μL*.

The basic dynamic processes include (i) invasion and depletion of RBC *x* by released merozoite (ii) parasite replication/growth via RBC invasion, and antigenic variation (switching rate, pattern); (iii) gametocyte conversion; (iv) immune stimulation and clearing of parasites by innate and specific immune effectors.

The model is run in discrete time steps, given by replication cycle (48 hrs for Pf). On each time step a new generation of merozoites {*r y*_*s*,*r*_} is released from the iRBC pool{*y*_*s*,*r*_}. They invade the available RBC *x* in proportion to their densities. Some of newly released merozoites can switch their var-gene expression. The AV switching process has two inputs, rate *ε* = 10^−4^ −10^−3^, and AV-pattern. Different switching patterns we proposed. We shall consider 3 of them based on loci ordering, (i) cyclic sequential (unidirectional) 1 → 2 → … → *m* → 1; (ii) symmetric sequential (bidirectional) 1 ↔ 2 ↔ … ↔ *m* ↔ 1; (iii) one-to-many (see [24]). In most simulations below, we use sequential patterns (i) and (ii).

#### Immune stimulation and clearing

All iRBC expressing a particular variant *r* can stimulate production of its specific immune effector (antibodies) *u*_*r*_, as function of their combined density 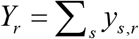. Also all of them are cleared by *u*_*r*_, depending on its level and specificity (affinity). The surviving iRBC fractions proceed to the next cycle, by releasing new generation of merozoites, and switching their var-genes (fraction *ε*). The details of immune stimulation – clearing (functions and parameters) are explained in the Supplement. Here we mention that immunity has finite memory, so IE variables {*u*_*r*_}decay gradually in time (c.f. [26], [38], [1]). This has implication for long-term (life-long) infection histories, a subject of future work. Further details on model variables, parameters, equations, inputs and outputs, are outlined in Supplement (Modeling setup).

#### Computer implementation and parameter choices

We set up and run our model for relatively small strain makeup (*m* = 5 − 10) var-genes, vs. realistic values 50-60 (c.f. [26]). One can think of such *m*, as an effective var-gene number. The *effective growth rates* of all variants were taken in the range, 4 ≤ *r* ≤ 8. The known Pf replication factors (merozoite /schizont) are typically higher (10-20). Our “effective” reduced *r*_*i*_ combine such replication factors with probability of successful RBC invasion. The combined growth rate *r*_*i*_ of a specific var-gene iRBC can be attributed to a combination of its *replication* and invasion potential of merozoites expressing this variant. Immune parameters (stimulation, clearing, waning rates) were chosen so that the resulting parasitemia patterns and RBC depletion in simulated histories, are overall compatible with malaria therapy data, though we didn’t attempt calibrating our system such data (c.f. [39], [40], [20], [41]).

All dynamic simulations, data processing and analysis, were implemented and run on Wolfram Mathematica platform. The essential feature of our model is dynamically varying system state (strain and immune-effector makeup), with new strains brought in by inoculae and old strains cleared out. Built-in Mathematica association tools allow an efficient implementation of such open-state systems.

## Results

### Single and mixed infection histories in naïve host

A typical naive infection history of a single 5-variant clone is shown in Figure 2. The panels include (a) RBC depletion reaching its peak (virulence = 20%) by day 37; (b) iRBC (parasitemia waves) with 5 expressed variants (thin curves) taking over in sequential pattern; (c) stimulated specific immune effectors; (d) gametocytemia; (e) probability of mosquito infection P(t).

**Figure 2:**
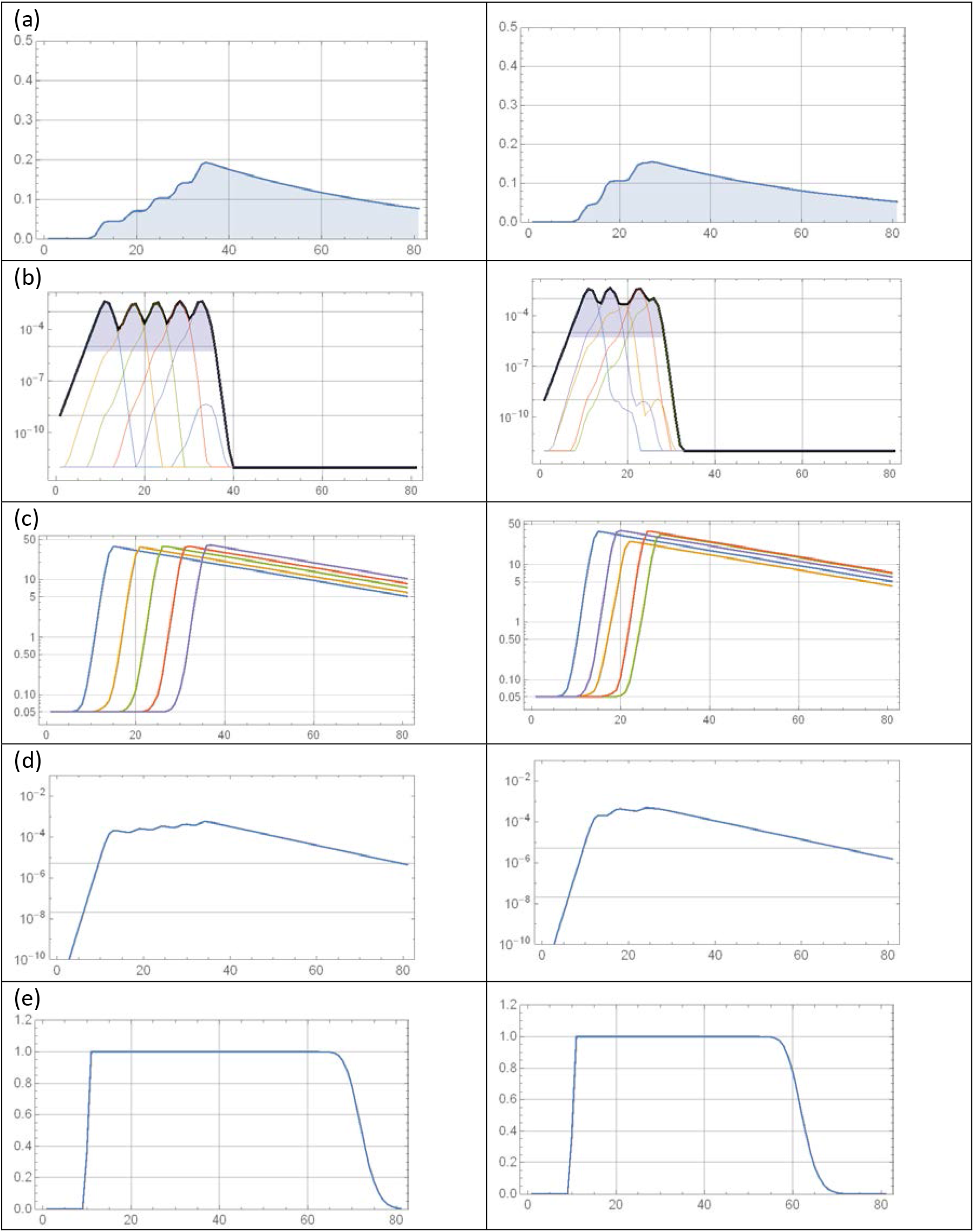
Naïve infection histories for 5V strain of clone (8, 4, 9, 13, 22). Two columns correspond to different AV switching patterns: (left) one-way sequential; (right) two-way sequential. Panel rows (top to bottom) represent a) RBC depletion; b) log-Parasitemia (iRBC) – total (solid black) and expressed variants (thin); c) variant-specific immune effectors; d) log-Gametocytemia; e) probability of mosquito infection. Two-way switching (right column) would shorten infection history, lower virulence (RBC depletion), and reduce transmissibility range.

The latter, P(t) is computed from gametocytemia curve G(t) panel (d), via a threshold-type sigmoid function *P* = *ϕ*(*G*), that predicts probability of mosquito infection in term circulating gametocyte level (see e.g. [42–44] and Supplement).

Function P(t) gives rise to an important strain ‘fitness output’ called *transmission potential* (TP). The latter is defined by integrating P(t) over a prescribed time window (e.g. infection duration [0, tf]), so TP = AUC (area under curve) of function P(t). Panel (e) shows TP is approximately equal to ‘host infectivity’ times ‘infective duration’.

Assuming uniform mosquito biting over time-range [0, tf], higher TP-value would imply higher likelihood of mosquito uptake and transmission; hence provide competitive advantage on population level.

We proceed to analyze naïve host histories, using different combinations of single and mixed-strain inoculae. Multiple factors can affect infection outcomes and estimated ‘phenotype values’: virulence (V), duration (D), TP. They include strain *immunogenic multiplicity d* = number of distinct var-genes (*d* ≤ *m* -number of loci), AV switching pattern, initial inoculum, and host immune status and competence.

For single-strain infections fitness traits (V, D, TP) are well defined, estimated from simulated histories in identical hosts. All 3 are correlated, higher V implies longer duration and higher TP. Initial dose has marginal effect, just shifting history by a few cycles. The AV-switching pattern had more pronounced effect (see Figure 2): symmetric sequential (ii) and one-to-many (iii) (not shown) give shorter D and lower TP, compared to direct sequential (i). Immune size *d* also matters, in general m-strain (number of var loci) of reduced diversity (*d* < *m*), behaves like d-strain.

The above ‘phenotypic’ strain classification breaks down for mixed infections, which exhibit more complicated dynamic patterns and multiple outcomes. Several factors can affect mixed infection histories, strain makeup, immune cross-reactivity (var-gene overlap), sequential order of shared var-genes, time lag between inoculae, et al.

We demonstrate it with a few examples of mixed infections and their TP-outcomes. Typically, mixed-strain competition would reduce individual TP (up to 50% or more), compared to uncontested (singe-strain) run (Figure 3, Table 1). But in some cases, we observe ‘cooperative behavior’, with some strains gaining TP, while others loosing (Supplement).

**Table 1:** TP outcomes for a 3-strain system (left column) in 3 contests: (i) Single (uncontested) shows near identical fitness score, (ii) synchronous initialization maintains TP2 (strain 2), but significantly reduces TP1 and TP3; (iii) delayed inoculum of #3 (by 4 cycles) vs. ##1-2, has marginal effect on TP2, lowers TP1 and drops dramatically TP3.

|  | Single | Sync | Delayed |
| --- | --- | --- | --- |
| { 10, 9, 19, 20, 1 } | 57.7 | 45.8 | 49.6 |
| { 7, 19, 0, 12, 14 } | 57.7 | 57.9 | 56.8 |
| { 3, 10, 16, 20, 9 } | 57.7 | 40.9 | 26.4 |

**Figure 3:**
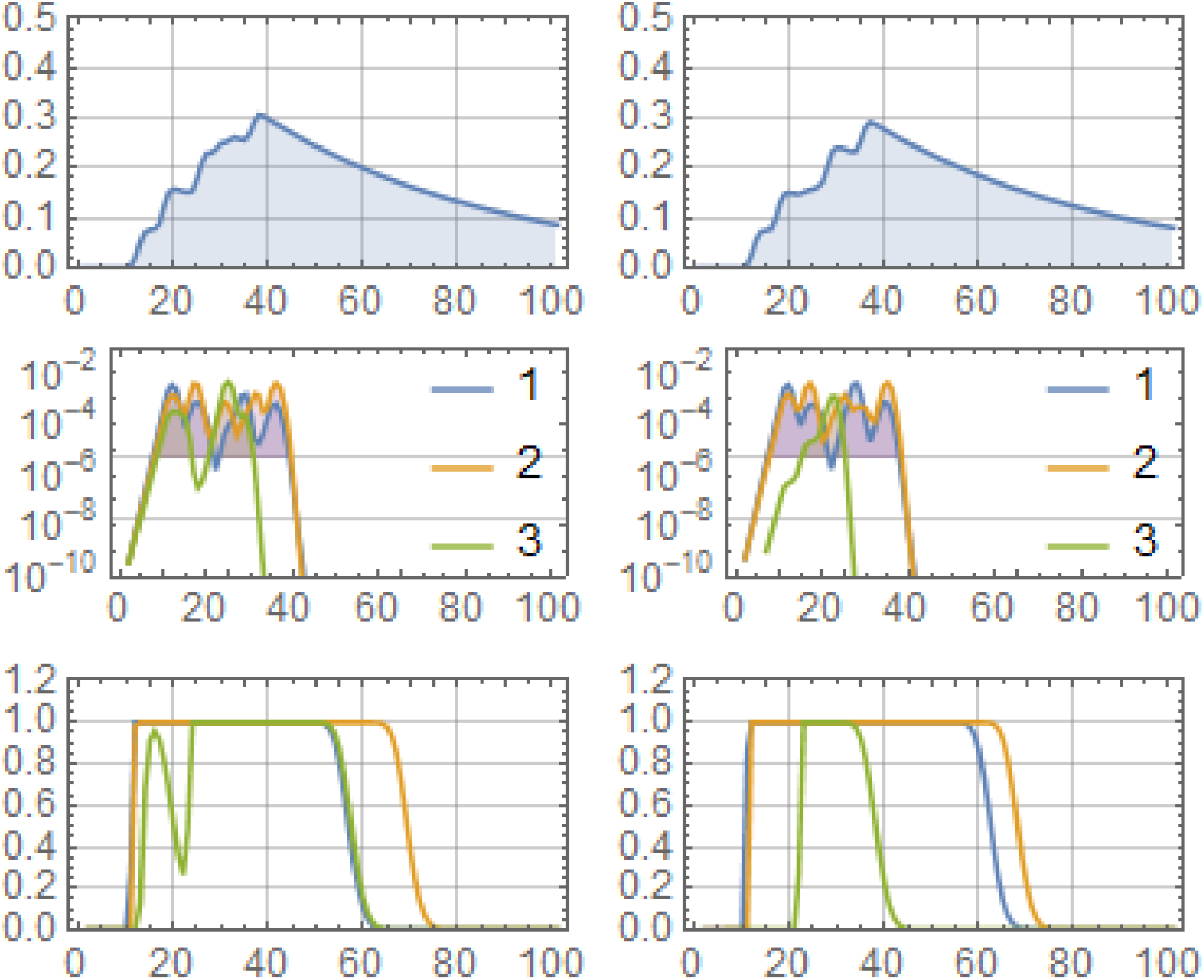
3-strain contest for triplet of Table 1. Panels in each column show (i) RBC depletion, (ii) strain parasitemia with MS-detectable level marked by shading, (iii) probability of mosquito infection. Left column has all 3 strains initialized at t=0; right column has #3 initialized 4 time units after ##1-2. It results in a marked drop of mosquito infection curve and its cumulative TP for strain #3.

### Immune-modulated parasite selection

Our main goal is evolution and selection of genetically structured malaria parasite in host populations. A realistic large-scale simulation would require structured host community (demographic, geographic distribution) coupled to mosquito environment. A community of human agents is coupled to ‘mosquito agents’ by randomly biting and transmitting circulating clones from ‘donor’ to ‘recipient’ host. Transmitted clones would undergo immune-regulated within-host process, as described above. In addition, they could recombine to generate new clones. Two processes, immune mediated ‘competition’ in humans, and ‘recombinant transmission’ in mosquito, have opposite effect on parasite diversity. The former tends to reduce it via immune clearing, the latter can potentially increase it. The two driving forces would shape parasite population structure and select ‘most viable’ type, or viable ‘cooperating cliques’ as we shall demonstrate below. To implement such fully coupled human-mosquito evolutionary system is challenging task that we shall address in the future work.

The current analysis is done under simplifying assumptions: (i) human population made of naïve hosts, (ii) no parasite recombination in mosquito. So mosquito part is limited to direct transmission of ‘infective’ parasite clones from donor to recipient. Both assumptions could be approximately valid under special conditions. For instance, naïve hosts can dominate transmission cycle in populations with high turnover rate relative to infection duration (so most of them experience a single bout of infection). Another example could be populations with the bulk of transmission carried by young children. Our reason for imposing the above assumptions is mostly technical (computational), rather than biological. The future work will extend the scope of modeling to more realistic host communities and environments.

The analysis of evolution and selection of multi-strain malaria will run in 2 steps: (i) *primary* selection in naïve host ensembles; (ii) *serial passage* (SP) in host lines and communities, over multiple transmission cycles.

### Fitness selection in naïve host ensembles

We want to identify most-fit parasite clones that could dominate in different patterns of mixed-infection. The model employs a hypothetical (‘natural’) var-gene pool, made of N=21 growth-types uniformly distributed over the range (4 ≤ *r* ≤ 8). We use it to generate a random repertoire of 200 clones {*s* = (*r*_1_, *r*_2_,…, *r*_5_)}, each one carrying 5 variants (cf. [26]). The 200 selected clones will comprise the entire parasite repertoire, as no mosquito-recombination is used in the current analysis. All var-genes {*r*_*i*_: *i* = 0,1,…, 20} are assumed immunogenically disjunct (no cross-reaction), but clones sharing identical variants could interact via immune regulation.

To select ‘most-viable’ clones from the repertoire, we run multiple simulations of single and mixed-infections with different strain combinations. The key *selectable trait* will be transmission potential (TP), introduced in the previous section.

Each clone from the repertoire is subjected to a range of single/mixed contests (double, triple etc.), and we estimate the resulting TP scores. The number of all possible contests (doublets, triplets et al) of 200 grows rapidly with the contest size (= 2, 3, …). So we won’t run all doublets or triplets, but limit our trial to representative ‘fair contests’, whereby each clone would compete against an equal number of randomly drawn contestants for each pool size (see Table 2).

**Table 2:**
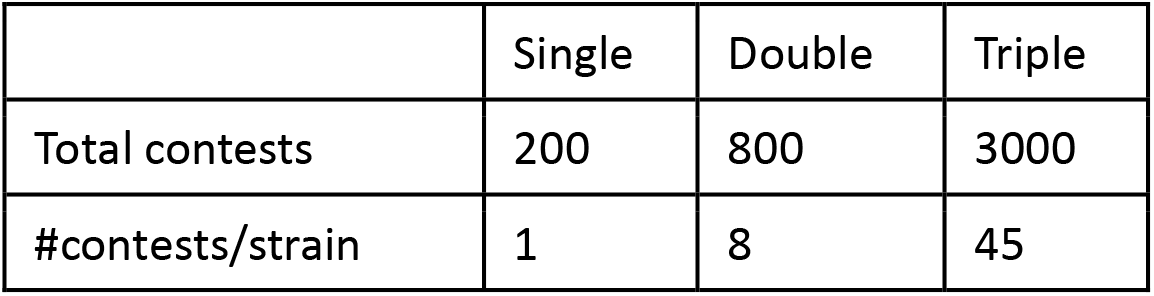
Fair contests for mixed infections (single, double, triple) for random strain repertoire of 200. In mixed cases (double, triple) each strain ‘plays’ against equal number of its partners (8 in double contest, and 45 in triple contest)

|  | Single | Double | Triple |
| --- | --- | --- | --- |
| Total contests | 200 | 800 | 3000 |
| #contests/strain | 1 | 8 | 45 |

TP-fitness is a well-defined trait for a single contest. But as we mentioned earlier, mixed-infections can give a range of TP-values, depending on the contest-pool makeup, host immune status, relative timing of inoculate et al (see Supplement). In such cases, we propose to define strain-TP by its mean value <TP > over all contests of a given type. Specifically, for each clone *s* we estimate its TP1-fitness (‘single contest’), TP2 –mean TP in double contests, TP3 – mean TP in triple contests (see Table 2).

Such mean <TP> gives relative success rate of strain *s* against multiple competitors drawn from large hypothetical test pool, where different mixtures are present in roughly equal proportions. Frequency of mosquito uptake of strain *s* in such system is proportional to its mean-TP, a combination of (TP1, TP2, TP3) weighted by fraction in the contest pool. So statistical selection of mixed-contest outcomes can identify clones most likely transmitted from *primary* (initial) host pool to the next generation.

For our statistical analysis we simulated 3 mixed contests (single, double, triple), to get for each clone its mean TP-values: TP1 (singlet), TP2 (8 doublets), TP3 (45 triplets). We comparted distributions of mean TP-values. The overall pattern was drop of <TP> with contest-size (single vs. double vs. triple). Figure 4 (a) shows distribution of *TP*_*n*_ for (n=1,2,3). A relatively narrow TP-range of singlets (natural TP-fitness), moves to lower and more broad range for doublets (about 15% drop), and further down for triples (20%). Panel (b) illustrates singlet-to-doublet loss by their scatter plot.

**Figure 4:**
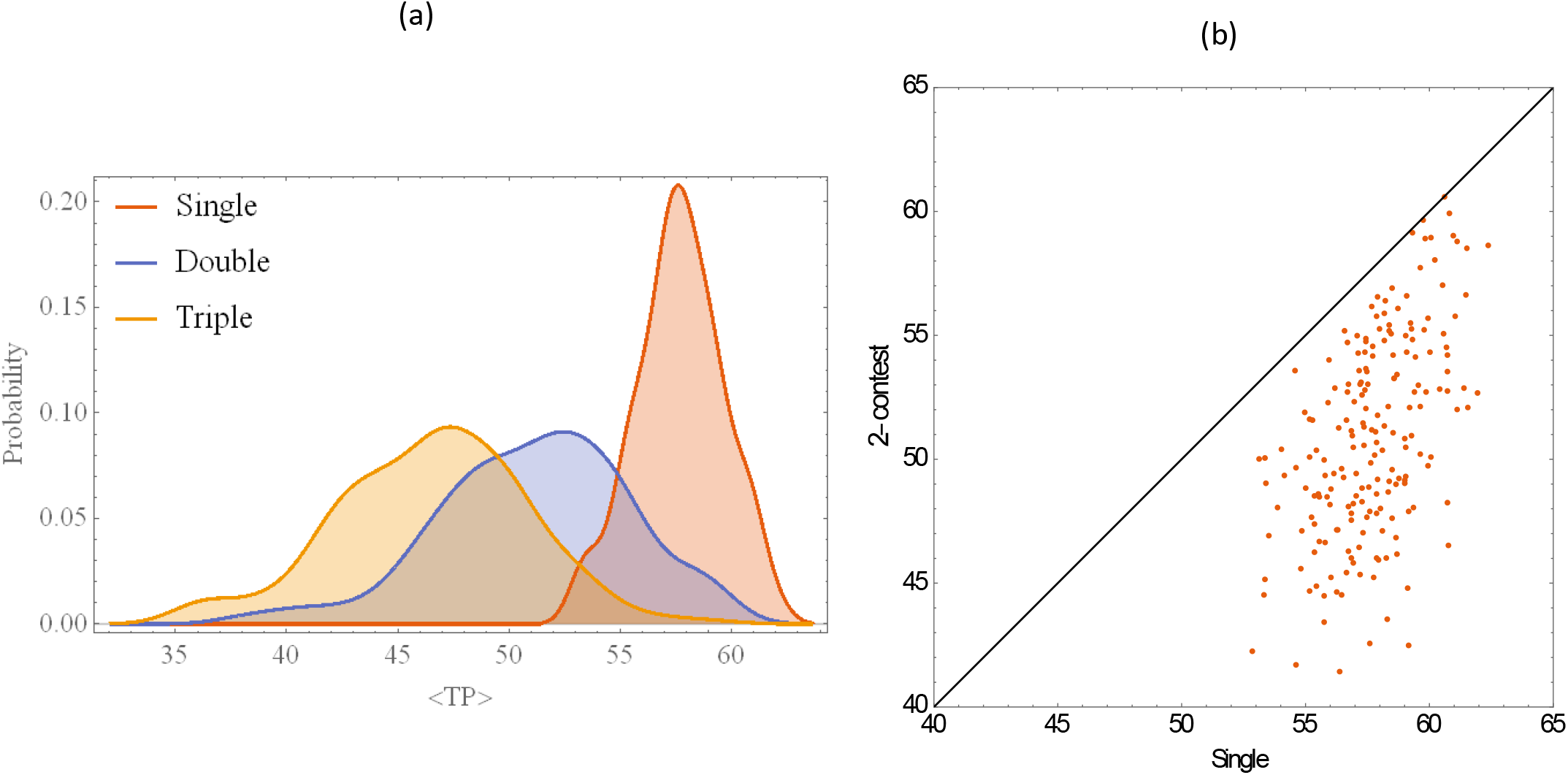
(a) Mean TP distribution for single, double, triple contest; (b) scatter plot of single TP vs. double-mean TP. The overall pattern is loss of TP-fitness in mixed competition

**Figure 5:**
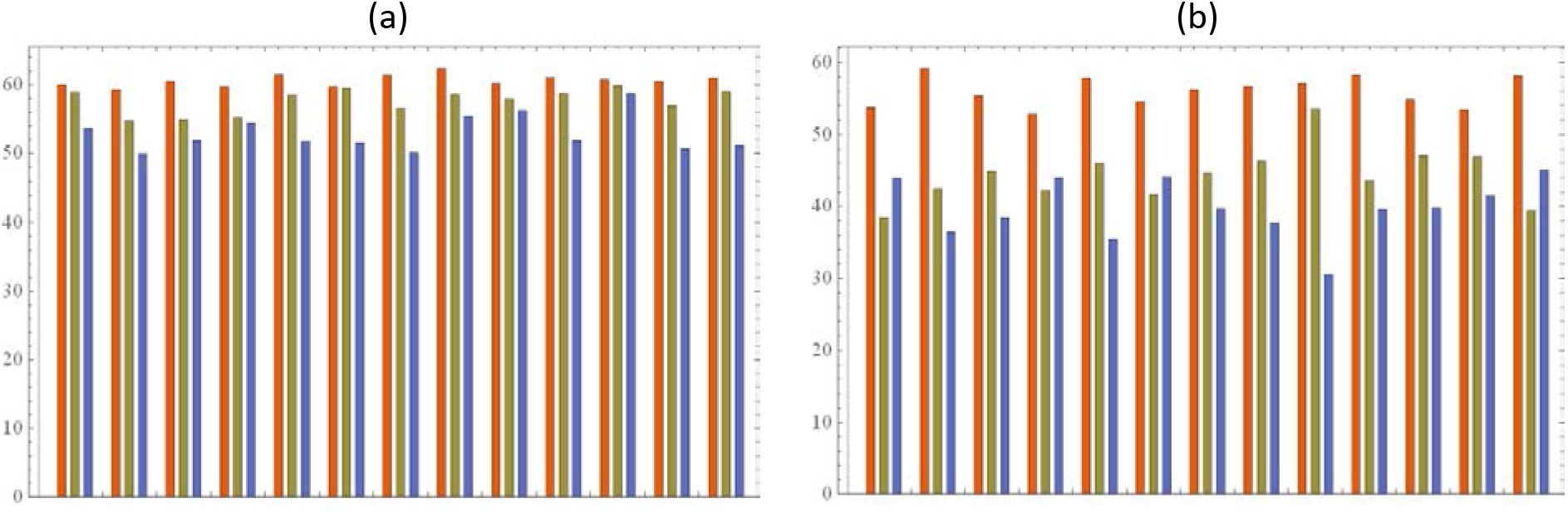
Mean TP drop for single (red), double (gray) triple (blue) contests. Panel (a) shows 13 best selected types (based on 80-th quantiles); panel (b) 13 lowest types.

Each contest (TP1-TP2-TP3) would select its own best-fit quantile clones. Their analysis reveals highly non-random primary selection. Indeed, the overlap of the 80% best quantiles of (TP12-TP2-TP3) contained 13 common clones, well above the expected (random) value 1.6. Furthermore, the best 13 types have shown only minor drop of their TP-values between (TP1-TP2-TP3). It suggests ‘highly transmissible’ types can maintain their dominance and TP-value, in multi mixed-strain contests (Figure 4(a)). In contrast, the bottom 20% quantiles are nearly random (overlap =2), and exhibit marked drop of the TP, from natural ‘singlet’ values TP1 to ‘doublet /triplet’ (TP2-TP3) (Figure 4(a)).

The selected 80% -quantile overlap was also found to correlate with high *mean-growth* phenotype (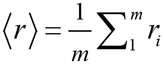), and high combined TP-score (TP1+ TP2+ TP3) (Supplement).

Besides clone selection, our mixed-contest runs have revealed highly non-random var-gene. To identify them, we assigned each variant *v* its frequency (multiplicity) in a given strain pool (# of clones containing *v*). It differs from the usual allele frequency count based on host population fractions. The original random repertoire of 200 was unbiased, with approximately equi-distributed var-gene frequency (mean = 47.6). The primary selection however, produced strong bias towards high-growth variants (see Figure 6). Therefore, mixed-contest immune competition produced highly non-random outcomes, both on the parasite level (selected clones), and in terms of their var-gene frequencies.

**Figure 6:**
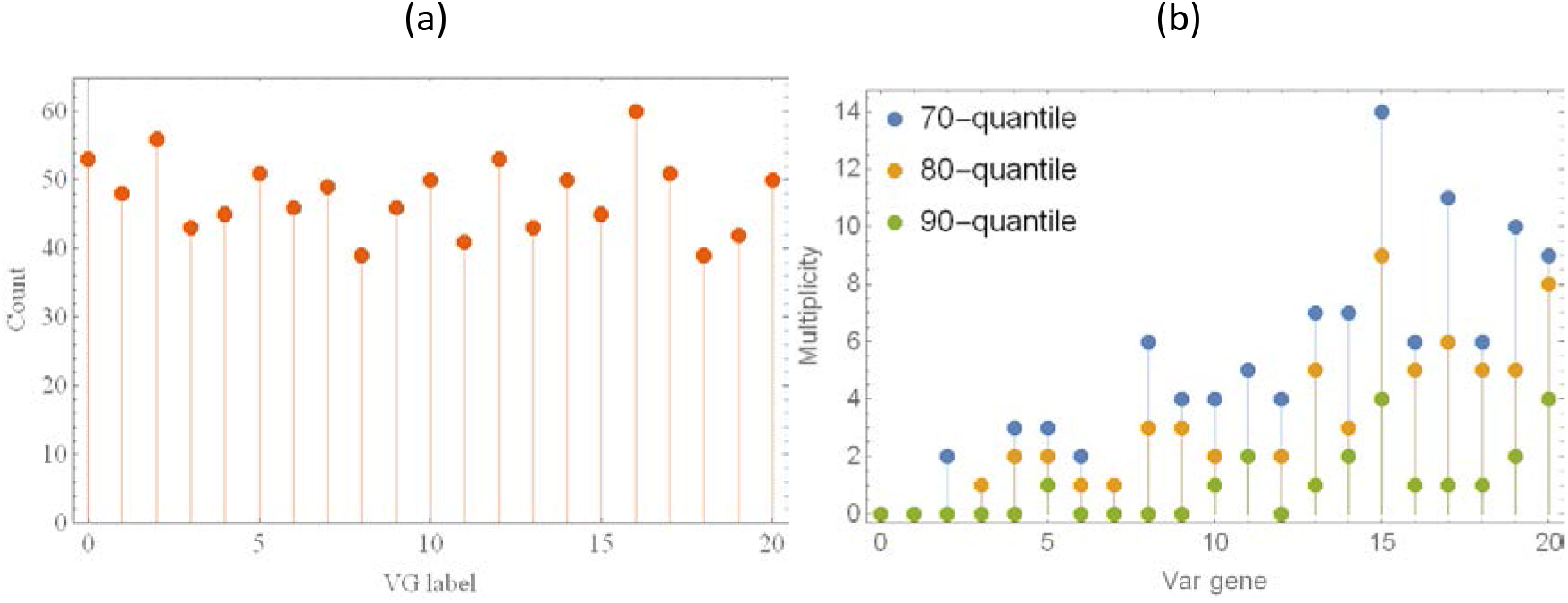
Var-gene multiplicity in the original random repertoire (a); and the best selected TP1-TP2-TP3 pool for high quantiles p=70-80-90%. High growth vars [10–20] dominate in contest selection.

One way to think of ‘primary selection’ is a hypothetical naïve host pool (e.g. children) invaded by a diverse set of mixed infections, drawn from large natural ‘parasite repertoire’. Regardless of initial distribution of such mixtures, the final output (transmissible clones) would be strongly biased toward winning types. Such clones are more likely to survive naïve (child) selection, and get transmitted to next host generations.

In the next section, we shall use *primary* selection *pool* to study evolutionary dynamics of multi-strain malaria over multiple transmission cycles and host generations.

### Serial-passage evolution in host lines and ensembles

A serial passage (SP) line follows a sequence of transmission cycles from donor to recipient. Each SP-agent runs its infection course from time 0 to *d*_*P*_ (passage day), then a collection of transmissible gametocytes (strain mixture) at *d*_*P*_ is transferred to the next in line. The passage day can be fixed, or vary randomly over a prescribed time window (*d*_0_ ≤ *d*_*P*_ ≤ *d*_2_). In our numeric experiments, passage day serves as proxy of transmission intensity – EIR (entomological inoculation rate), shorter *d*_*P*_ meaning higher EIR. In most experiments below we use random *d*_*P*_.

Parasite mixtures undergo selective processes, as they follow through SP-lines and multiple transmission cycles, whereby successful types or groups take over. While individual SP-lines may differ, statically significant results arise on population level, i.e. ensembles of SP-lines run in parallel.

Evolutionary time scale used in our analysis, requires some clarification. Transmission cycles serve as discreet time units, as each line proceeds over generations (1^st^, 2^nd^ …). Though different SP-lines of an ensemble may not be synchronous in real physical time (due to random *d*_*P*_), a fixed generation of all SP-lines can be viewed as (approximate) ‘instantaneous’ snapshot of the entire community.

We are interested in statistical evolution of parasite population structure in such SP host ensembles. More specifically, we look at persistence and frequency distribution of single and mixed parasite clusters, the resulting relationship (network structure) among selected clones, their var-gene makeup and expressed phenotypes.

The analysis makes a few simplifying assumptions on transmission dynamics. We drop Intermediate parasite stages and processes in human and mosquito, so ‘transmissible gametocyte pool’ from donor turns into recipient ‘merozoites/iRBC inoculum’. The net effect of omitted stages and processes are time lags in the onset of infection, which are negligible on evolutionary time scale (generations). As above, all hosts in SP-lines are assumed naïve.

We conducted two types of SP experiments using parasite repertoire of the previous section. In the first experiment, the 95% best quantile of primary selection was used to initialize all SP-lines in the host ensemble. No recombination or external inputs were allowed, so initial mixture could only get depleted in such evolutionary process.

In the second experiment, we allowed an external input on each SP-step, drawn randomly from the complete parasite repertoire (an additional mosquito bite). This case allows parasite diversity to increase and sample more broadly the entire 200-clone repertoire.

In both cases, we study long-term evolution of parasite population structure, the resulting statistical and dynamic patterns, including stable dominant strains and mixtures, their genetic makeup, phenotypic profiles et al.

#### SP evolution for confined parasite pool

The first set of SP-experiments employs 10 best-fit strains of the primary selection (Table 3) for initialization. Each SP-lines was run over multiple cycles, and we generated evolving host community of 200 SP-lines. To study the effect of transmission intensity we used 2 time windows for random passage day: (i) *d*_*P*_ = 15 − 30 (high intensity), (ii) *d*_*P*_ = 25 − 40 (low intensity).

**Table 3:** Selected 95% quantile of the TP1-TP2-TP3 primary contest

|  |  |  |  |  |  |
| --- | --- | --- | --- | --- | --- |
| 1 | 18 | 5 | 10 | 16 | 15 |
| 2 | 17 | 13 | 20 | 10 | 7 |
| 3 | 18 | 13 | 11 | 16 | 4 |
| 4 | 15 | 20 | 19 | 17 | 11 |
| 5 | 13 | 16 | 20 | 18 | 0 |
| 6 | 20 | 12 | 13 | 15 | 5 |
| 7 | 17 | 20 | 15 | 3 | 4 |
| 8 | 19 | 16 | 0 | 20 | 10 |
| 9 | 20 | 18 | 13 | 19 | 3 |
| 10 | 20 | 13 | 9 | 17 | 10 |

A typical SP-line history with high intensity random *d*_*P*_, is shown in Figure 7. On each cycle we measured strain PT (probability of mosquito transmission) at the passage day. It varied in the course of history, with most strains maintaining highest level 1, while other (#7) getting depleted after 30 cycle. Figure 8 shows evolution of two SP-communities with high and low transmission intensity. Each curve shows host population fraction, carrying a particular strain mixture. In both cases (i-ii) evolution drives such system to fixation. High intensity case (i) maintained elevated strain diversity (only one clone, #7 got depleted). Low intensity case (ii) splits into several small (4-and 5-strain) persisting clusters, with mean diversity dropping from 10 to 6.

**Figure 7:**
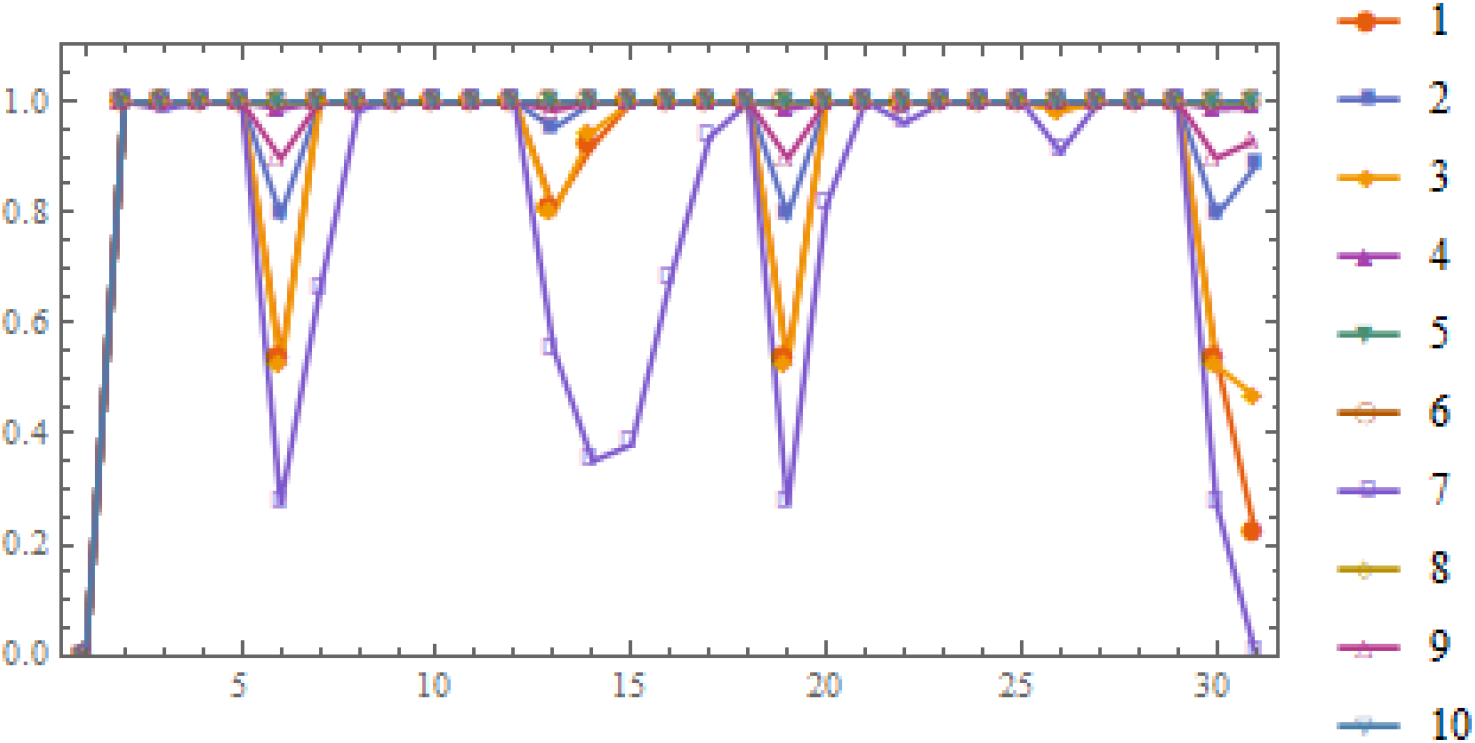
A typical SP-line of 10-starin system (Table 3) over 30 cycles, with random *d_P_* = 15 − 30. On each cycle we measure strain PT (probability of mosquito transmission) by the end of cycle. Most strains persist over the entire span, but some (#7) get depleted by the end (30^th^ cycle)

**Figure 8:**
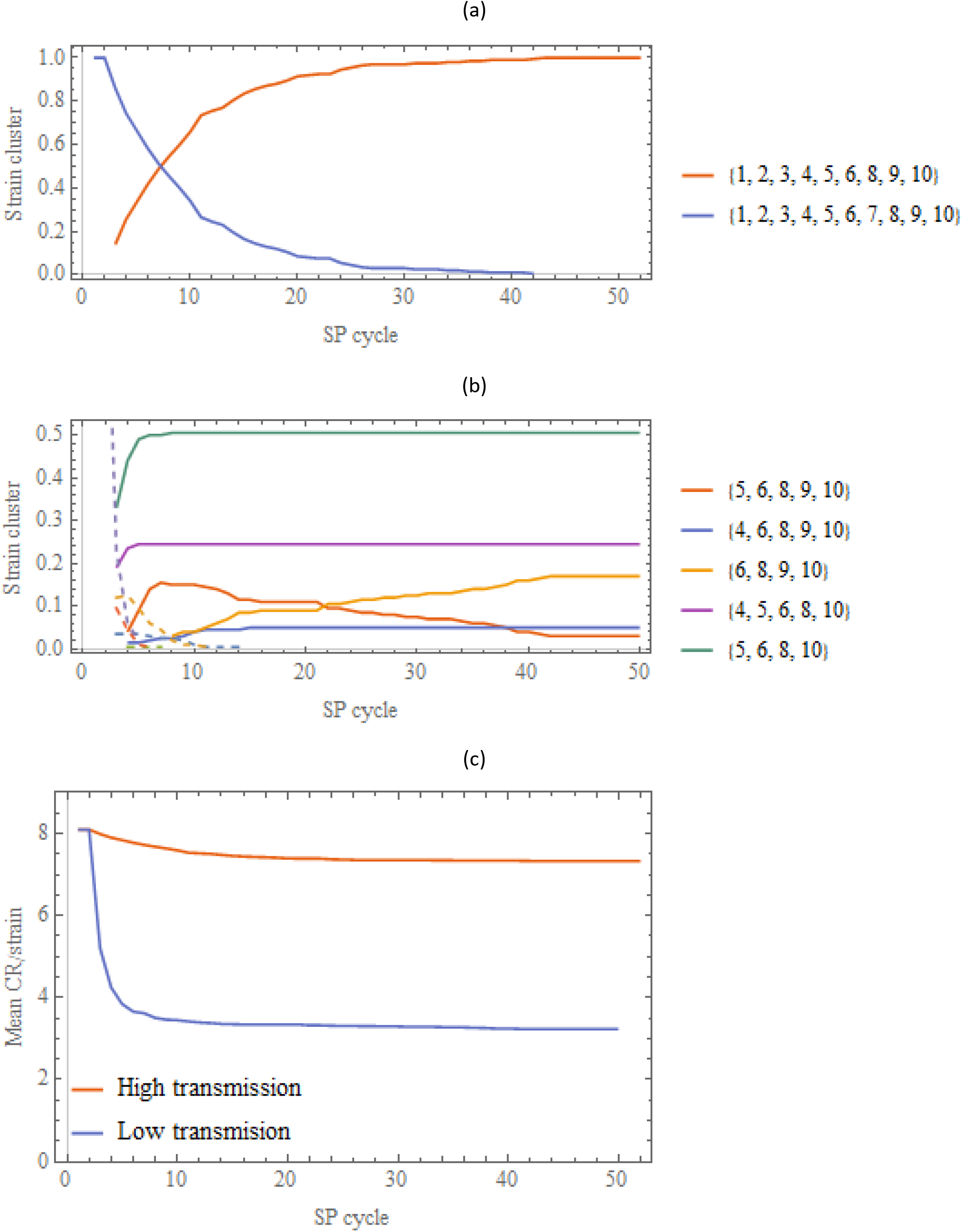
Long term evolution (frequencies of strain clusters) in SP-ensembles initialized with 10-strain mixture (Table 3) and different intensity of transmission: (a) *d_P_* = 15 − 30; (b) *d_P_* = 25 − 40. Panel (c) shows evolution of mean CR/strain in 2 cases

##### Strain organization and connectivity

There different ways to describe strain patterns in terms of their structure and dynamics. A typical dynamic evolution produces long term persistent groups of ‘cooperating’ strains (see e.g. Figure 8). Structurally, they consist of overlapping clusters; each cluster in turn can be described by its immunogenic profile, represented by cross-reactivity (CR) matrix or graph.

We define CR-graph of strain complex {*a*, *b*,…} (nodes), by linking clones that share common var-genes; each link *a* ↔ *b* is weighted by the number (*C*_*ab*_) of shared variants. Figure 9 illustrates the CR-network of ‘best-fit’ pool of Table 3. This network is tightly bound, with weights varying from 0 to 4 (e.g. *C*_1↔9_ = 1, *C*_2↔10_ = 4). V*ertex degree C*_*a*_ is weighted network is defined, as the sum of all link weights (Σ_*b*_*C*_*ab*_), connected to *a*. Mean cross-reactivity of cluster {*a*, *b*,…} is defined as total CR (Σ_*a*_*C*_*a*_) divided by the cluster size. For the primary selection pool of Table 3, mean CR = 8.

**Figure 9:**
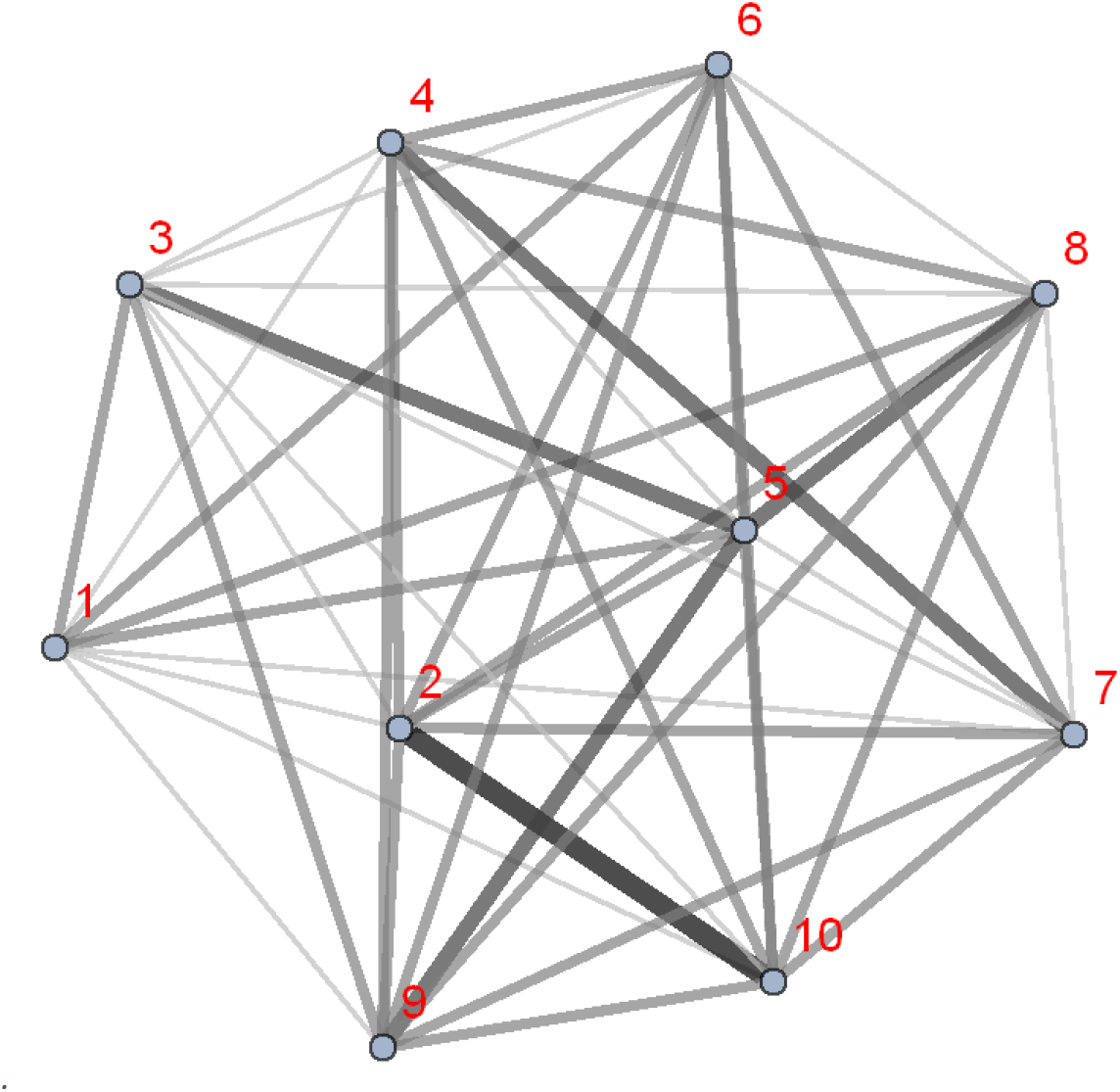
Tightly linked CR network of 10 selected clones (Table 3). Link weights (1,2,…) are marked by thickness and shade of gray, e.g. weight (1-9) =1, weight (2-10) = 4 (4 shared var genes)

The loss of parasite diversity in SP-evolution is expected to lower its mean CR. Figure 8 (c) shows the CR-drop in both cases (i-ii). Such CR-loss is qualitatively consistent with theoretical predictions of strain theory, whereby immune selection should drive parasite distribution towards disjunct clusters (see e.g. ([13–15]).

More close analysis of the terminal evolution state (Figure 8(b)) shows two-level organization into persistent interconnected clusters. The three dominant clusters of Figure 8(b), CL1= (5, 6,8,10), CL2=(4,5,6,8,10), CL3=(6,8,9,10) share common core (6,8,10) (Figure 10). Theirs CR-graphs (Figure 10 (b) show reduced connectivity, compared to strongly linked initial system (Figure 9).

**Figure 10:**
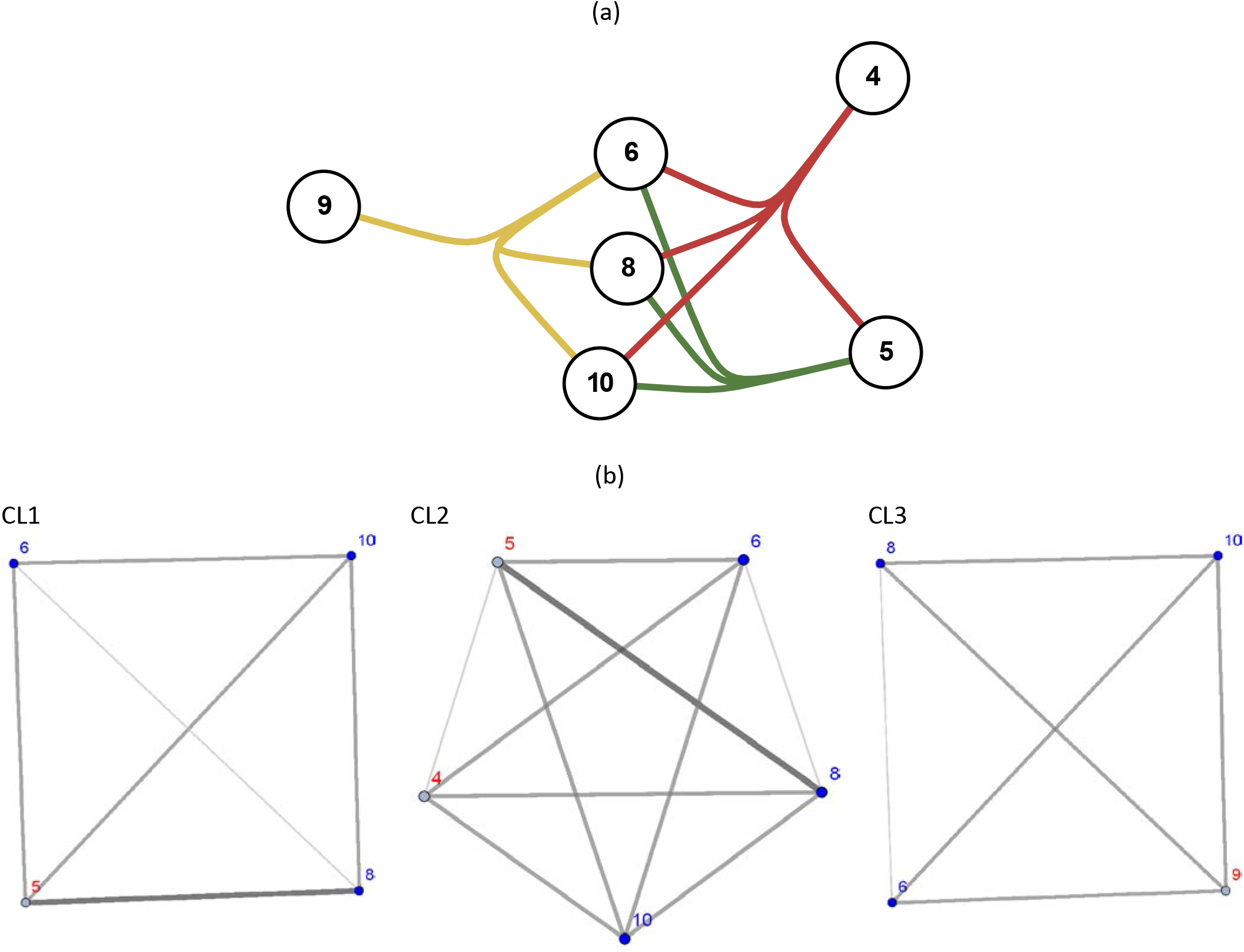
(a) Three dominant clusters of Figure 8(b): CL1= (5, 6,8,10), CL2=(4,5,6,8,10), CL3=(6,8,9,10), share common core ##6,8,10. (b) their CR networks; the core nodes marked in blue

Table 4 summarized mean CR-connectivity of core-strains (6,8,10) in different configurations: complete 6-node terminal network of Figure 10 (column “Total”), followed by 3 dominant clusters (CL1, CL2, CL3), and the overlap (core) triplet (6,8,10). We observe a consistent drop of mean CR. It suggests evolution of the initial 10-strain system towards lower CR-connectivity state, consistent with general principles of immune selection in mixed infections, and the strain theory (c.f. [1, 5, 24, 26]).

**Table 4:** Mean ‘CR/node’ for 3 core strains ##(6,8,10) in the complete residual complex ##(4,5,9,6,8,10) of Figure 10; 3 dominant clusters, and the core triplet (6,8,10)

|  | Total | CL3 | CL2 | CL1 | Core |
| --- | --- | --- | --- | --- | --- |
| 6 | 1.6 | 1.3 | 1.4 | 1.3 | 1. |
| 8 | 1.6 | 1.3 | 1.6 | 1.5 | 1. |
| 10 | 1.8 | 1.5 | 1.6 | 1.5 | 1.3 |

#### SP-evolution with external source

Our second experiment involves the entire 200 repertoire of the previous section (primary selection). Here we use as an external source of inoculae, rather than initialization, which will play secondary role. So a serial passage will combine transmissible donor output, as before, supplemented with randomly drawn clone from the entire 200-repertoire. Such community can be viewed as subpopulation (children) submerged into a larger community and transmission environment with multiple circulating strains.

As above, our goal is to study evolution of parasite population structure in multiple SP-lines and ensembles. The analysis of dynamic patterns becomes more involved now, compared to the previous (limited-pool) case, as many more persistent clones and ‘cooperative cliques’ can arise here.

We start with strain-diversity analysis, then move to identifying ‘persistent clones’ and ‘cooperative cliques’, their organization (strain clustering, CR-connectivity, var-gene makeup).

As above, we run simulations of multiple SP-lines, over long evolutionary path (100 to 200 generations), with two types, high (15-20) and low (25-40) transmission intensity. Figure 11 (a-b) shows typical SP-histories for high/low intensity cases. Multiple clones from the entire 200-pool enter these histories, but they exhibit vastly different duration patterns, varying from few cycles to long evolutionary spans. Their temporal distribution is strain-specific and highly non-random. The core-nodes of the previous selection (##6-8-10) persisted throughout the entire period (they correspond to 3 long dark lines at the bottom of panel (a)). But new persistent clones arise, generated by random source, which were not high-quantile winners of ‘primary selection’.

**Figure 11:**
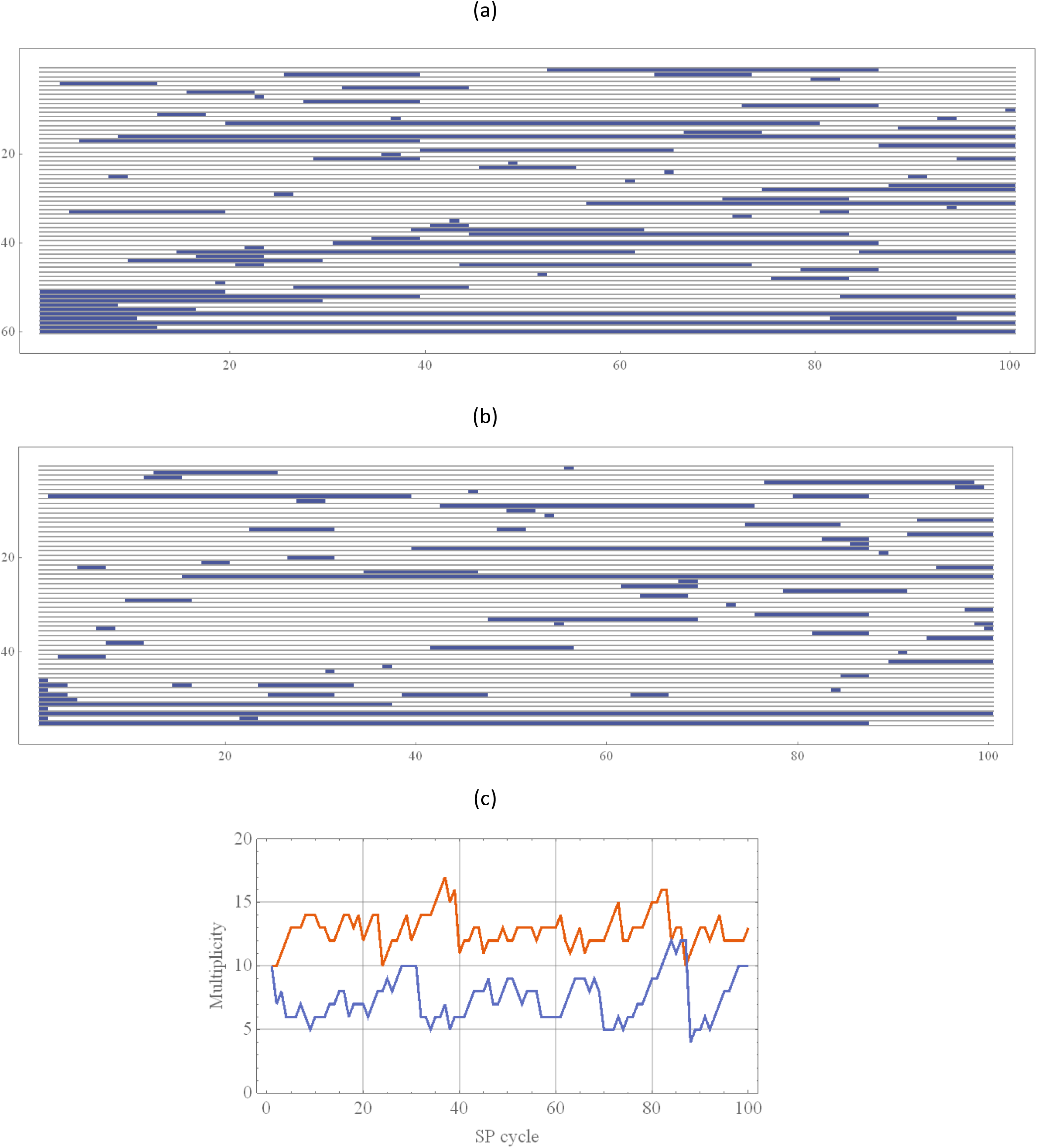
Typical SP –lines of 100 generations with random inoculae in 2 cases: (a) high intensity *d_P_* = 15 − 30, (b) low intensity *d_P_* = 15 − 30. Each row marks a particular clone appearance and duration (shaded). History (a) contained 60 clones altogether, while (b) had 55 (reduced diversity). The clone ordering in (a)-(b) is not the original 200-repertoire, so identical numbers could be different types, except the low left corner (initial inoculum) made up of the original 10 best-fit selection of the previous section. Panel (c) shows multiplicity (strain diversity) of histories (a)-(b).

We are interested in statistically stable patterns, i.e. ‘persisting strains’ and statistically significance clusters in the entire SP-ensemble. We define ‘statistical significance’ as ensemble frequency above 25%-threshold, and call the resulting persistent clusters *cooperative cliques*. In the previous (limited-pool) case, core nodes ##6-8-10, played this role.

Two types of SP-ensembles, ‘limited-pool’ and ‘open-source’, exhibit markedly different dynamic patterns and statistical behavior. The former case had all SP-lines going to fixation. The latter case does not go to ‘dynamic fixation’ (as new clones are brought in), but it can produce stable statistical patterns, in terms of ensemble frequencies. We first ask for a statistical fixation (population frequency) of individual clones. Figure 12 shows evolution and selection of such ‘persistent clones’, defined by host frequency (count) above 25% of population, at the late stage of evolution (t > 150).

**Figure 12:**
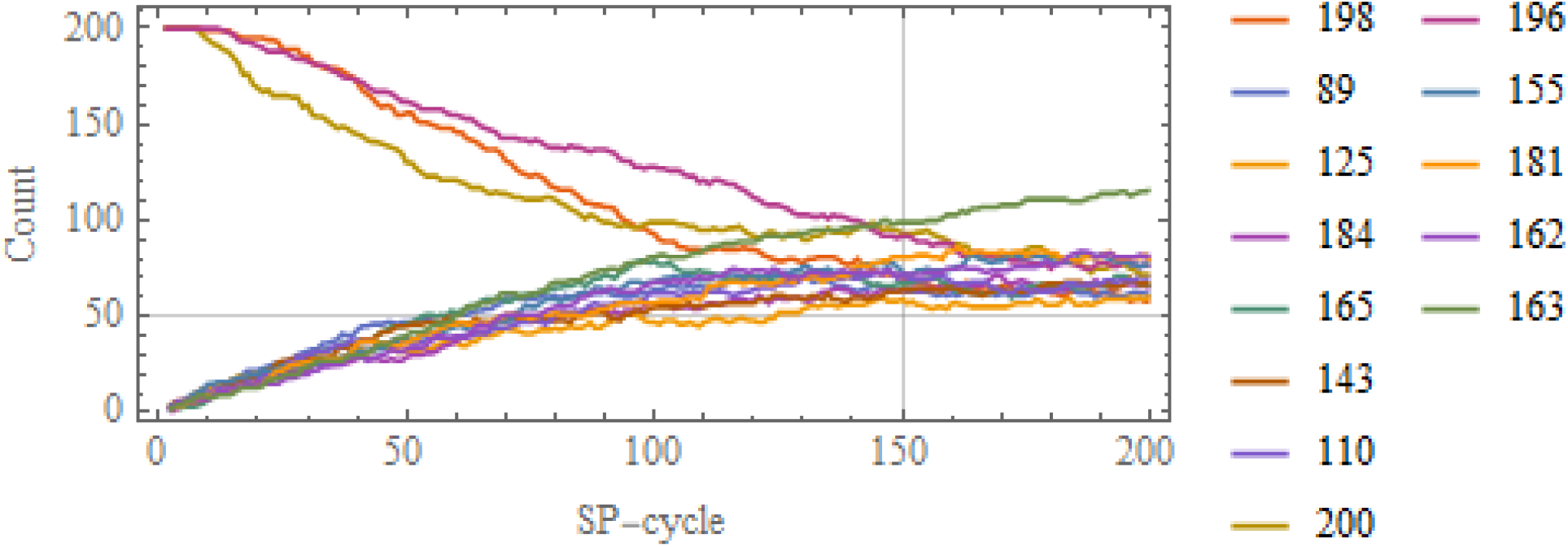
Evolutionary path of high-frequency clones of the SP-ensemble of 200 lines. Each curves shows population frequency of selected clones (number of SP-lines that carry the clone). Selection was based on frequency > 50/200, attained after 150 generations.

Altogether, we observed 14 such ‘persistent clones’, but only three of them (##196-198-200) came from original primary-quantile pool, used for initialization. As expected, those 3 were the core-triplet of the previous ‘limited pool’ case, labelled there as ##6-8-10. The old core-triplet (6-8-10) was now joined by 11 newly acquired clones from the random source.

Individual-clone selection of Figure 12 however, gives only partial view of SP-ensemble evolution. Beyond individual persistent strains, it also exhibits a cooperative behavior. But this time *cooperative cliques* (singlet, doublet, triplet, etc.) are defined statistically, via ensemble frequency persisting over sufficiently long period [*t*_0_, *t*_1_] of evolutionary history.

To extract such cliques, we took an ensemble of 200 SP-lines over 200 transmission cycles, with random passage date, 15 ≤ *d*_*P*_ ≤ 30. The time window for clique selection was chosen as 50 < *t* < 150. In this time-window, we observed 196 persistent combinations with the following distribution of multiplicities

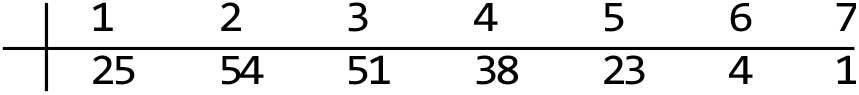

i.e. 25 persistent singlets, 54 – doublets, etc. The most crowded clique had 7 cooperating clones.

Further examination of persistent cliques has shown many of them sharing the same limited pool of *core nodes* (Table 5). It appears successful clones are able to collaborate with multiple partners in different combinations.

**Table 5:** 11 common-core clones shared by double-triple-quadruple,-quintuple cooperative clusters in simulated SP-ensemble with open source. In each row, number on the left is clone’s label in the 200 repertoire, the remaining 5 digits give its var-gene makeup

|  |  |  |  |  |  |
| --- | --- | --- | --- | --- | --- |
| 32 | 20 | 13 | 9 | 17 | 10 |
| 44 | 19 | 13 | 1 | 10 | 2 |
| 68 | 19 | 16 | 0 | 20 | 10 |
| 105 | 20 | 10 | 1 | 15 | 9 |
| 124 | 18 | 14 | 4 | 1 | 8 |
| 132 | 19 | 2 | 8 | 11 | 3 |
| 158 | 19 | 1 | 17 | 0 | 10 |
| 172 | 20 | 12 | 13 | 15 | 5 |
| 179 | 20 | 2 | 5 | 18 | 7 |
| 180 | 19 | 12 | 1 | 16 | 10 |
| 184 | 20 | 0 | 8 | 15 | 10 |

Figure 13 (a) shows the resulting CR-network of the core nodes, and panel (b) shows their population frequencies in the SP-ensemble, over time-window [50-150]. Some of them, ##32-68-172, attained high frequency values: (60-70) of 200 host lines. It means large host-population fractions of the SP - community (30-35%) carried 3 dominant clones over long evolutionary history.

**Figure 13:**
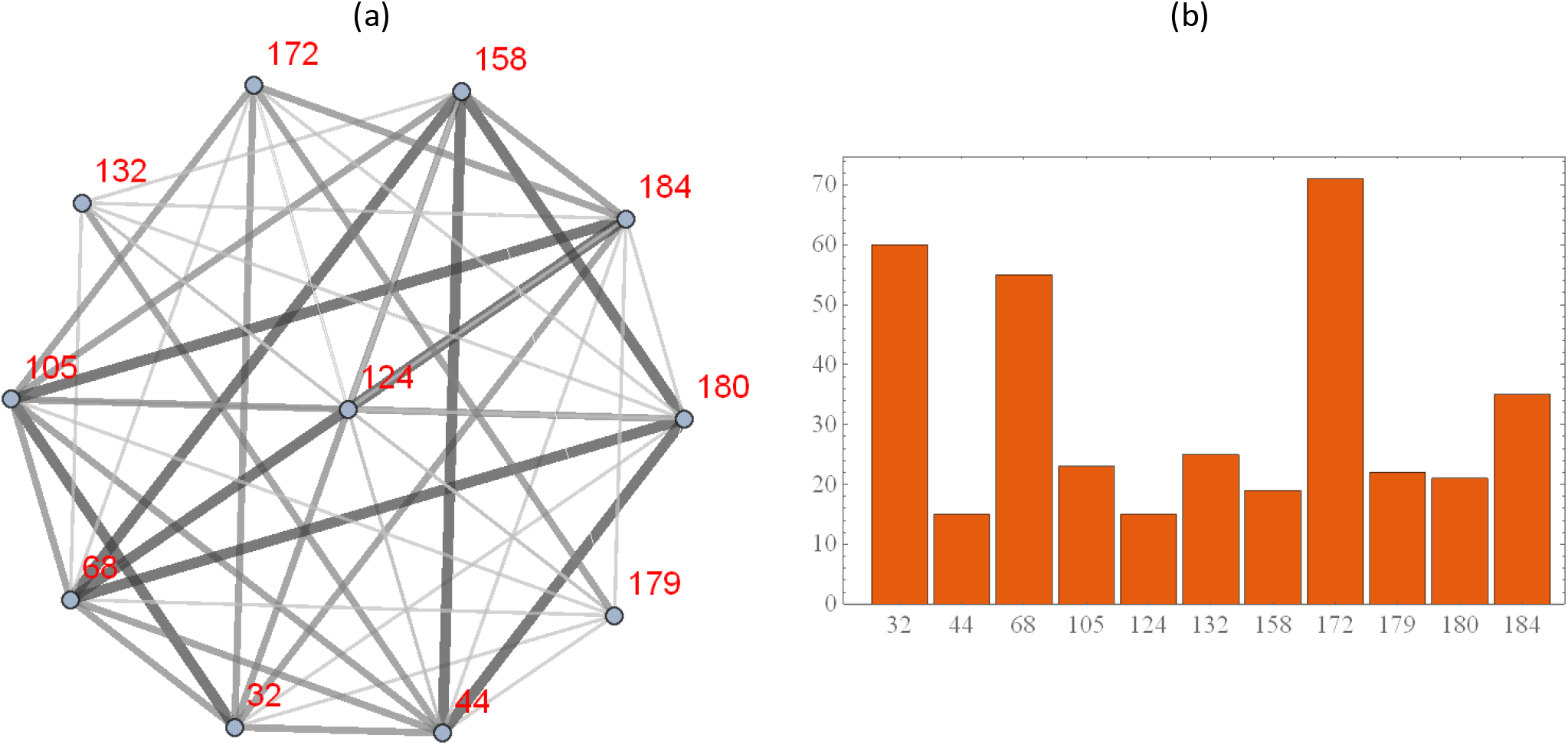
(a) CR network of the core-strain selection of Table 5; (b) their frequencies in the SP-ensemble of 200 lines over the time-range [50-150].

**Figure 14:**
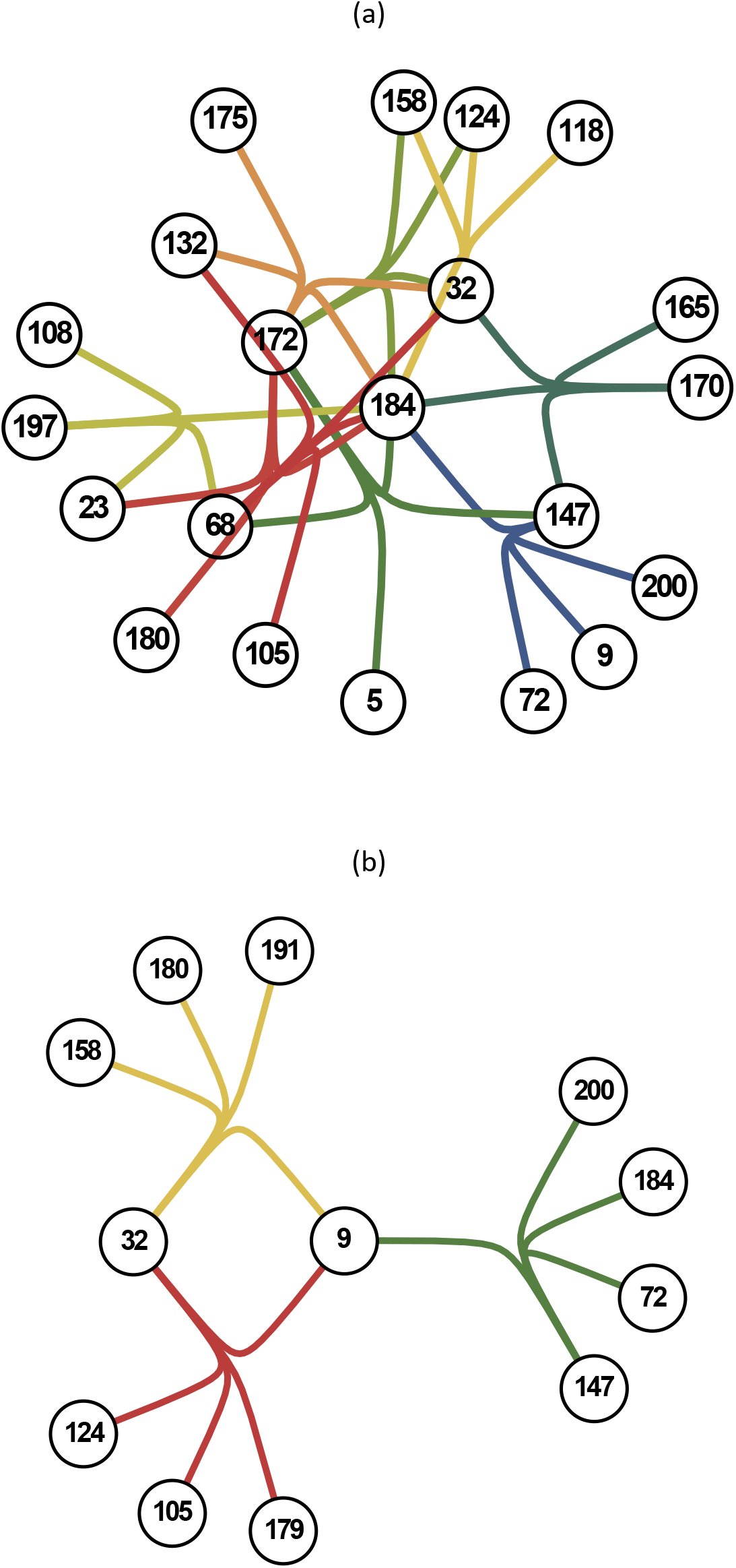
Persistent 5-cliques of the core node # 184 (a), vs. non-core #9 (b). In either case, the clique-graph consists of 4 nodes (tree-like structure) headed by #184 or #9.

Overall, 11 cooperative clones exhibited relatively high CR-connectivity (Figure 13 (a)), and some peculiar patterns of their genetic makeup (Table 5). Namely, locus 1 was occupied by high-growth variants (##18-20) of the var-gene pool, while the remaining loci contained middle or low growth var-genes. Individually (single-clone infections), such strains would produce high-growth initial wave of parasitemia, and relatively high virulence (RBC-depletion)

We also found the best core-nodes of the open-source evolution (Table 5) differ from the previous (limited-pool) case (Table 3).

Each core-node of Table 5 carried different (low and high multiplicity) cliques, including doublets, triplets, et al. The persistent cliques of the open-source evolution demonstrated stable *cooperative patterns*, similar to *stable clusters* of the previous (limited-pool) case. We show (Figure 15) persistent 5-cliques of the *core-node* (#184), and compare it to *non-core node* (#9). In both cases, the resulting networks consist of several associated tree-like graphs, headed by #184 or #9, with linked branches (shared strains). The highly connected ‘cooperative’ node #184 has 9 branching clusters, interlinked by the fellow core-members (##32, 68, 132, 124, 158, 172) (Table 5), each core carrying its own linked cluster. This can be contrasted to a pair of disconnected cliques of #9, with a single CR-link produced by the core node #32.

Overall, we find the open-source SP-evolution leads to formation of long-lasting network, made of a few dominant *cooperating nodes* with high CR-linkage, and multiple weekly-linked tree-like clusters attached to the dominant nodes. Qualitatively, it resembles some observed serum-dominance networks [6], and network patterns produced in numeric simulations of [1, 5].

## Comments, conclusions

### Immune selection vs. transmission intensity

The key drivers of parasite dynamics and evolution in host populations, immune clearing and transmission intensity, have opposite effect. The former, according to so-called ‘strain theory’ (see e.g. [13–15, 24]) is expected to drive cross-reacting parasite clones apart into disjunct clusters (host subpopulations), hence reduced CR. The latter has opposite effect, by maintaining multiple strains together, or expanding them, via horizontal mosquito mixing, hence increased CR. The net balance depends on relative strength of ‘immune segregation’ vs. ‘transmission spread / mixing’. Two examples of SP-evolution for confined strain pool demonstrated the effect of such balance. High transmission rate limited immune effects by maintaining diverse strain pool and high CR-connectivity. Low intensity produced greater strain segregation, clustering with reduced diversity and lower CR. The immune effect was limited in the former case, due to shorter infection duration, while the latter allowed more persistent ‘segregation’. We also observe organized network structures that arise under ‘weak mixing’ and ‘strong segregation’, dominated by few core nodes, and weakly linked clusters attached to the core.

The open-source SP allowed us to sample the complete strain repertoire. It produced markedly different selection patterns, where ‘cooperative’ behavior proved more important, than ‘competitive fitness’.

### Limitations

The present study was confined to naïve hosts. It includes individual histories, host ensembles, SP lines, et al. The ‘naïve-host’ assumption can be justified under special conditions, e.g. relatively short host turnover compared to infection duration (so each host can experience at most a single episode), or the bulk of transmission carried over by naïve hosts (children). Neither one is particularly realistic. Here we adopted ‘naive-host’ hypothesis for technical reasons, and the future work will extend the scope of model and analysis to fully coupled systems.

Another important component to be developed is a proper account of innate immunity in malaria control, the present models confines it to febrile control.

### Future model development; control implications

The next step in model development and applications will include long-term (life-long) infection histories for demographically structured host populations, and multiple transmission cycles. Furthermore, one can add parasite recombination in the system, during mosquito sexual reproduction stage, as well as ectopic (haploid) recombination within-host. In such systems parasite diversity is no constrained by any prescribed strain pool, but can evolve depending on transmission intensity, host makeup (immune status) and possible interventions (drug treatment, ITN). Next step would to set up spatially (geographically) distributed host communities, and mosquito environment. Similar to [35], one can explore links between mosquito distribution and transmission intensity and parasite diversity (multiplicity of infection) for individual hosts, risk groups and host communities.

Another possible application of multi-strain models is to track geographic spread of infection in distributed host communities, using clones as biomarkers.

There are many ways to expand our model and make it consistent with the available molecular datasets on var-gene makeup and diversity (e.g. [5]). Such calibrated (validated) multi-strain systems could have potential applications for malaria control.

One example could be a putative PfEMP1-based vaccine to prevent severe (cerebral) malaria in children, (see e.g. [45]). Many forms of severe malaria are associated with cyto-adherent phenotype of iRBC, controlled by var-genes. Such cyto-adherence leads to formation of iRBC complexes glued together (rosettes) that clog small capillaries. Naturally acquired immunity against severe malaria is associated with disruption of rosette formation, hence a putative anti-PfEMP1 (var-gene) vaccine preventing cyto-adherence. Such vaccine could also lower parasitemia levels within-host, and inhibit its transmission in host communities.

An apparent obstacle to the concept of PfEMP1-vaccine is natural diversity of var-gene pool in host populations [2, 5, 32]. Our analysis suggests ‘dominant types’ of selected parasite strains and their var-genes could be more limited. However, it is not clear, whether such ‘selection’ could provide a practical choice for vaccine targets.

An alternative approach to PfEMP1-vaccine would be to extend the current modeling setup, where each var-gene is treated as a ‘single immunogenic unit’, to more detailed ‘structured’ var-gene system, with sub-units containing multiple (variable, conserved) domains that could provide suitable vaccine targets (see e.g. [33]).

## Supporting information

Supplement - modeling setup

## Acknowledgement

Our modeling project on malaria immunology was initiated during DG sabbatical leave at NIMBioS (University of Tennessee, Knoxville). We thank L.J. Gross and V. Ganusov for hospitality, and S. Karl for productive discussions on the subject.

## Supplement

### Modeling setup

In-host modeling of malaria was subject of many studies, see e.g. (^1–5^, ^6^,^7–10^,^11^). Our setup shares some features of papers (^12^, ^13–15^), but the details, and computer implementation differ. We outline it below.

#### Genetically structured parasite

Parasite strains are described as combinations of var-genes, *s* = (*r*_1_,…, *r*_*m*_), each var-gene having its prescribed growth/replication factor (*r* = *r*_*i*_), and specific antibody *u*_*r*_. The in-host system consists of 3 cell populations (per muL of blood): *x* – target RBC, **y** = (*y*_*s*,*i*_)-infected (iRBC) of strain s, expressing EMP variant *r*_*i*_, gametocytemia **G** = (*G*_*s*_), and immune effector variables **u** = (*u*_1_, *u*_2_,…), labeled by var-genes. The system’s state changes in discrete time steps, dt = 2 days (merozoite replication cycle):

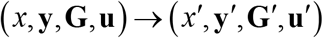

### Variables and equations

**Target** (susceptible) RBC population *x* is subject to homeostatic maintenance, survival (with 100-day life span), and depletion by invading merozoites. Newly invaded RBC pool by merozoite population *M*, and susceptible RBC *x*, is given by *Y* = (1 − *e*^−*pM*/*x*^)*x*, *p* – probability of invasion, or 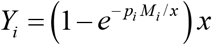 for multi-strain system (*Y*_1_;…;*Y*_*n*_).

**Parasitized** iRBC, **y** = (*y*_,*i*_) release merozoites *M*_*s*,*i*_ = *r*_*i*_*y*_*s*,*i*_, with replication factors (*r*_*i*_), determined by their var-genes. At the start of each 2-day cycle these merozoites invade susceptible RBC, and after AV-switch, they create a new generation of iRBC (*Y*_*s*,*i*_). On each cycle a fraction of iRBC 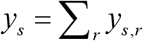 is converted to **gametocytes** *G*_*s*_, that circulate in blood (have life-span 8-10 days), and mature. The mature fraction (= 20% of G) can be uptaken by mosquito.

#### Antigenic variation (AV)

Different types of AV switching mechanism and matrix *A* = (*ω*_*ij*_), 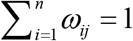, are used in the literature: uniform, sequential, random, structured e.g. “one-to-many” (see ^2,5 11,12,16,17 15 18^). In our setup, we adopted a sequential (band) matrix

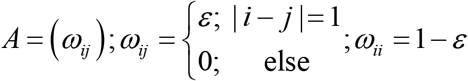

whereby a fraction of newly released merozoites of strain *s* = (*r*_1_,…, *r*_*m*_), can switch its expressed type, *M*_*s*,*i*_ → *M*_*s*, *j*_ to adjacent loci (*j* = *i* ±1), at a prescribed rate *ε*.

#### Innate and adaptive immune clearing

*Innate* (febrile) control works as density-dependent switch (sigmoid) function,

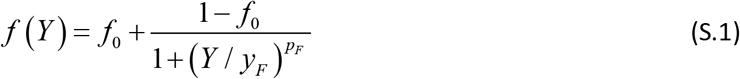

of total newly-released parasitemia (iRBC) *Y* = Σ_*s,i*_ *Y*_*s*,*i*_. It can suppress (clear) all iRBC, up to a fraction (*y*_*s*_ → *f*_0_ *y*_*s*_), at high total parasitemia *Y*. But febrile control is switched off at low parasitemia.

*Adaptive immunity* clear specific parasite iRBC strains expressing variant *r* by factor *ϕ* (*K u*_*r*_), depending effector variable *u*_*r*_ and affinity *K*. Clearing function *ϕ* (*Ku*) is derived in below (equations (S.6))

Specific IE (Ab) *u*_*r*_ are stimulated by circulating iRBC (*y*_*s*,*r*_) expressing variant *r*. They in turn, bind to such iRBC, and clear them depending on their concentration (value *u*_*r*_). Immune effectors (*u*_*r*_) also maintain their baseline level, and have finite memory (life-span 50-200 days), similar to RBC.

**Immune stimulation** of specific effector *u*_*r*_ is determined by its antigenic trigger *Y* = *Y*_*r*_, in our context all iRBC (circulating strains {*s*}) that express variant *r*, *Y*_*r*_ = Σ_*s*_*y*_*s*,*r*_. Variable *Y* stimulates proliferation of immune B-cells, their conversion to secreting (plasma) cells and release of antibodies (effector *u*). We lump all these processes into a single immune stimulation function,

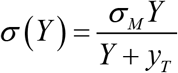

with maximal proliferation rate *σ*_*M*_, and sensitivity threshold *y*_*T*_. So *σ* (*Y*)changes from a ‘decay mode’ (*σ* (*Y*)< 1) at low trigger (Y) to the ‘growth mode’ (*σ* (*Y*)> 1), as *Y* passes a critical threshold *y*_*T*_. Exponential growth phase of immune effector *u* cannot proceed indefinitely, but it should slow-down at high *u*-values (increased B-cell population). In our model we implement such constraining density-dependent factor, 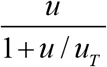. So out model would allow near-exponential growth (proportional to *u*) for low values (*u* < *u*_*T*_, below threshold *u*_*T*_), but for high *u*, the growth rate turns into a constant, determined 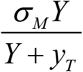. Hence, immune-stimulation function takes the form

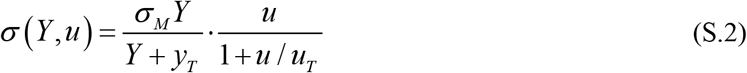

We also allow for slow waning of immune effector *u* at a certain rate 0 < *δ*_*u*_ < 1, due to antibody degradation and memory loss.

#### Equations

To simplify notations we label expressed parasite clones by index *i* = 1, 2,…; each label *i* = (*s*, *r*) combines ‘strain’ *s*, and expressed variant *r*; AV-switching matrix *A* = (*ω*_*ij*_)

**Table 1:**
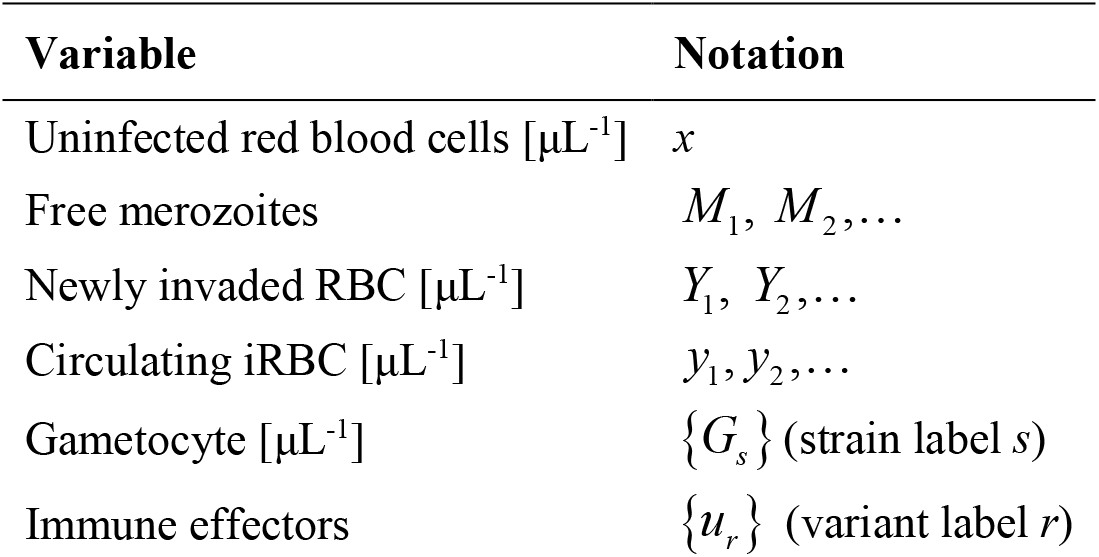
Dynamic variables and auxiliary functions

**Table 2:**
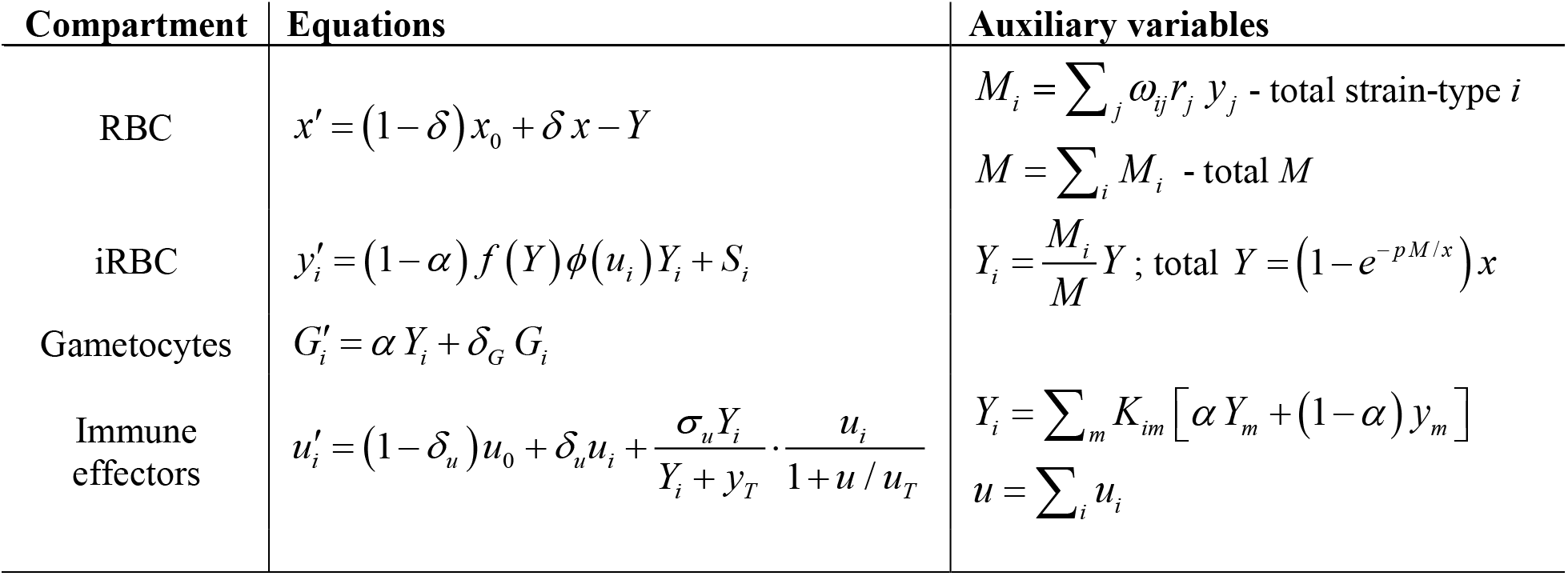
Model equations (single-strain case)

| Compartment | Equations | Auxiliary variables |
| --- | --- | --- |
| RBC | $x' = (1 - \delta) x_0 + \delta x - Y$ | $M_i = \sum_j \omega_{ij} r_j y_j$ - total strain-type $i$<br>$M = \sum_i M_i$ - total $M$ |
| iRBC | $y'_i = (1 - \alpha) f(Y) \phi(u_i) Y_i + S_i$ | $Y_i = \frac{M_i}{M} Y$ ; total $Y = (1 - e^{-pM/x}) x$ |
| Gametocytes | $G'_i = \alpha Y_i + \delta_G G_i$ | |
| Immune effectors | $u'_i = (1 - \delta_u) u_0 + \delta_u u_i + \frac{\sigma_u Y_i}{Y_i + y_T} \cdot \frac{u_i}{1 + u / u_T}$ | $Y_i = \sum_m K_{im} [\alpha Y_m + (1 - \alpha) y_m]$<br>$u = \sum_i u_i$ |

**Table 3:** Parameter names and values/ranges used in the model

| Parameter Definition | Notation | Value range |
| --- | --- | --- |
| Normal red blood cell level [$\mu\text{L}^{-1}$] | $x_0$ | $5 \times 10^6$ |
| Red blood cell loss rate (survival fraction)<sup>1</sup> | $\delta$ | 0.98 |
| Parasite clearing cut off [$\mu\text{L}^{-1}$] | $y_C$ | $10^{-6}$ |
| Parasite replication factors [per 2-day cycle] | $r_i$ | 4-8 |
| AV-switch rate [per cycle] | $\varepsilon$ | $10^{-4} - 10^{-5}$ |
| Gametocyte conversion fraction | $\alpha$ | 0.001-0.005 |
| Innate (febrile) clearing rate (max) | $f_0$ | 0.2 |
| Febrile threshold (iRBC/ $\mu\text{L}^{-1}$) | $y_F$ | $0.01 x_0$ |
| Adaptive immune stimulation threshold (iRBC/ $\mu\text{L}^{-1}$) | $y_T$ | $(0.001 - 0.003) x_0$ |
| Baseline immune level (dimensionless) | $u_0$ | .05-.1 |
| Immune efficiency (cross-reactive affinity) | $K_{ij}$ | $\{0,1\}$ |
| Saturation threshold for immune stimulation/growth | $u_T$ | 20-50 |
| Maximal proliferation rates | $\sigma_u$ | 2-4 |
| Immune memory (loss rate/cycle)<sup>2</sup> | $\delta_u$ | 0.9 - 0.95 |

### Kinetics of immune clearing

Anti-malaria immunity involves different types of antigens and immune effectors (Abs). An important, part of it are anti-erythrocyte EMP immune effectors (*u*_1_;*u*_2_;…). They bind to respective var-antigens expressed on iRBC *y*_*i*_, or other cross-reacting types. We want to estimate the surviving fraction *ϕ* (*u*) of newly invaded iRBC *Y* (by the end of merozoite cycle), as function of immune effector *u*. Derivation below employs continuous Ab-binding (opsonisation) process described by differential equations (DE), but we use terminal values of the resulting solution functions at *T* = 48 hrs.

To get the surviving fraction of IRBC by the end of cycle (*T*) one can proceed in two ways. Both involve binding-dissociation kinetics of antibodies *u* to iRBC *y*. In the first case, this process is coupled to subsequent removal of opsonised iRBC at a prescribed clearing rate *ν*. In the second case, we work with opsonisation fraction of iRBC, represented by function 0 < *s* (*t*)< 1, and assume hypothetical *survival probability* function of such opsonised iRBC. We outline and compare both approaches

#### Binding-dissociation kinetics

We call free and bound iRBC populations *y*(*t*) *z*(*t*), functions of the continuous time *t*. The schematic view of such reaction system

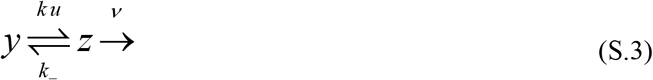

We take binding rate, *ku*, proportional to IE-concentration function *u*(*t*), and fix dissociation rate *k*_−_. Opsonized population *z*(*t*) is assumed to be cleared (by monocytes, spleen) at a fixed rate *ν*, i.e. removed by the end of the merozoite cycle (no progeny). So only free (unbound) population *y* is left to release merozoites in the next clonal cycle. We want to estimate the surviving fraction of initial iRBC population *Y* by the end of the cycle 0 < *t* < *T* (= 48 hrs). The DE system for reaction kinetics (S.3),

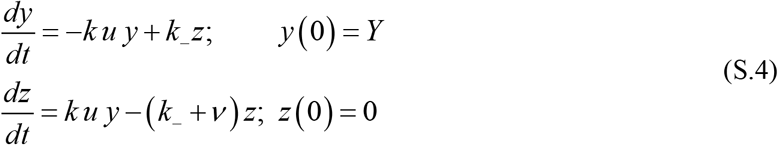

initialized at values *y*(0) = *Y* = (1 − *e*^−*pm*/*x*^)*x*, *z* (0)= 0, all newly invaded iRBC unbound by *u*. After *rescaling* over dissociation rate *k*_−_, *k* → *K* = *k*/*k*_−_ -affinity, *v* →*ν*/*k*_−_, assuming near constant (mean) effector level *u* over the cycle, we write solution of (S.4) via eigenvalues of matrix

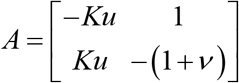

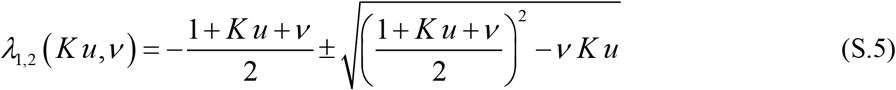

The resulting survival fraction is given by

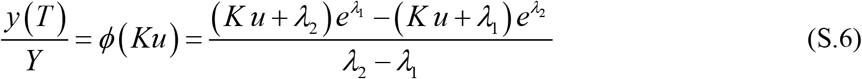

Function *ϕ*(*Ku*) approximately predicts surviving fraction of iRBC by the end of cycle in terms of dimensionless *Ku*, and relative clearing rate *ν*=*ν*/*k*_−_.

#### iRBC opsonisation

Here the basic dynamic variable is opsonised fraction, 0 < *s* (*t*)< 1, viewed as “mean iRBC coverage by specific Abs *u*”, or as an approximate fraction of “opsonized iRBC”, slated for removal. The corresponding binding-dissociation kinetics is given by DE

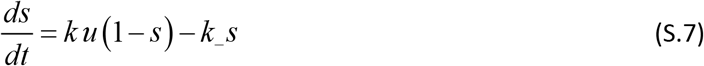

Next, we assume survival probability of opsonized iRBC is given by a sigmoid function, 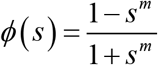 with exponent *m* (Figure 1). At high Ab-level *u*(*t*), solutions (S.7) rapidly approach equilibrium value 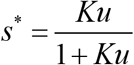. The resulting survival probability of iRBC *Y* is estimated, by factor *ϕ*(*s*^*^(*u*)). Two derivations produce somewhat different functional form of immune clearing (surviving fraction of *Y*). But qualitatively they like similar (Figure 2), and have similar effect on discrete-time dynamics of infection histories.

**Figure 1:**
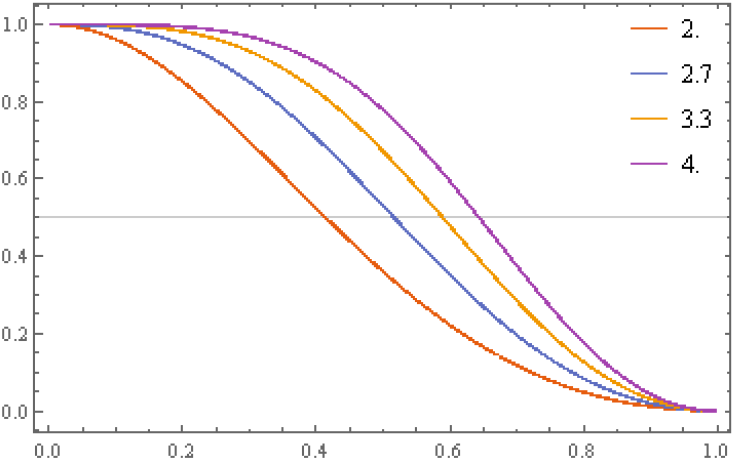
Survival function *ϕ*(*s*) for fractional coverage (opsonisation) 0 < *s* < 1, for several values of exponent 2 ≤ *m* ≤ 4

**Figure 2:**
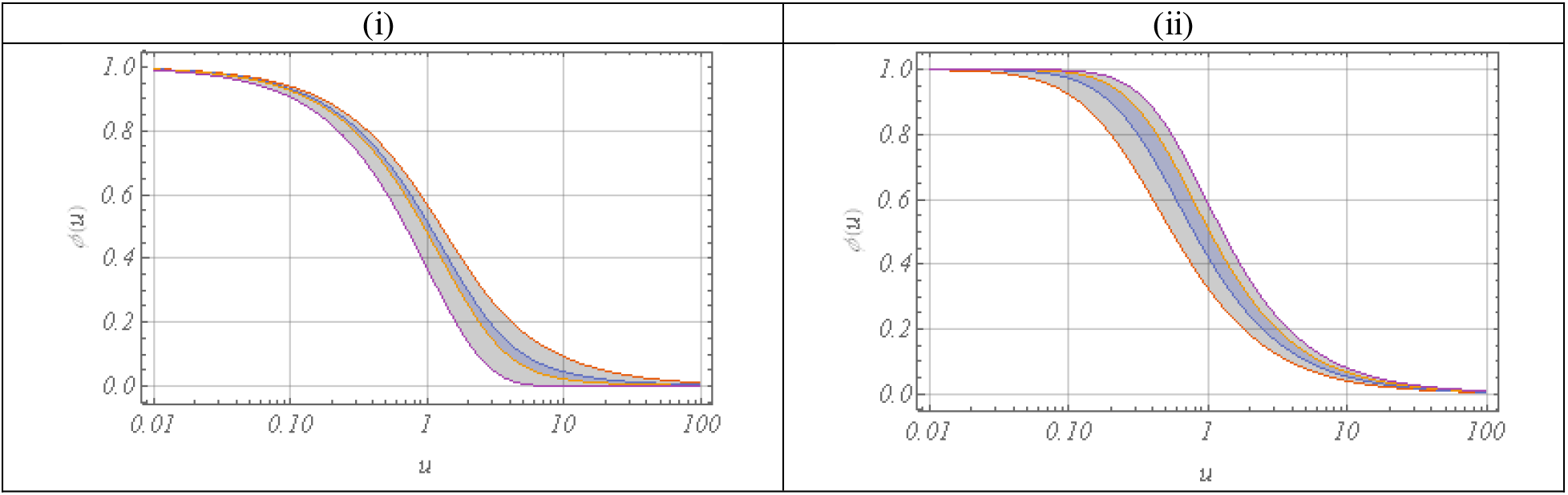
Survival iRBC-fraction *ϕ*(*u*), as function of immune effector *u*, in 2 cases: (i) binding-dissociation kinetics for a broad range of clearing rate (0.1 <*ν*< 10), top-to-bottom bounds of shaded envelop; (ii) opsonisation system for a range ‘clearing exponent’ 2 ≤ *m* ≤ 4

## Footnotes

1 Based on 100-day life-span of RBC

2 Based on immune (Ab) duration of 40-100 days

