## Supplement - modeling setup for "Competition, cooperation and immune selection of multi-strain Plasmodium falciparum malaria"

### Supplement: in-host multi-strain malaria system

D. Gurarie

#### Model setup

**Target cells** (RBC): susceptible population *x*; homeostatic maintenance, depletion by invading merozoites.
**Parasites**: iRBC $y$ (circulating and sequestered), $M = r\,y$ merozoites, r-replication factor

**Immune effectors** [1], [2] (specific Ab): anti-erythrocyte *u* (binds to EMP, prevents sequestration); anti-merozoite *v* (binds to MSP, prevents RBC invasion).

Antigenic variation (AV): each strain consists of m EMP-variants that stimulate variant-specific effectors $(u_1;\ldots;u_m)$, and a single (non-specific) MSP – effector v

#### Processes

**Invasion**: newly invaded RBC $Y = \left(1-e^{-pM/x}\right)x$, *p* – probability of invasion, or $Y_i = \left(1-e^{-p_i M_i/x}\right)x$ for multi-strain $(Y_1;\ldots;Y_n)$

**Immune clearing**

Innate (febrile) control "switch" $f(TY) = \dfrac{1}{1+\left(TY/y_F\right)^p}$ - density-dependent surviving fraction of I ( $TY = \sum_i y_i$ - total parasitemia iRBC), $y_F$ - febrile threshold

Adaptive effector variables (Ab) u

- Anti-erythrocyte: $\phi(K\,u)Y$ – surviving fraction of newly invaded RBC *I* (u- effector level, K – affinity). For cross-reacting multi-strain systems $\phi\left(\sum_j K_{ij}\,u_j\right)Y_i$. Clearing function $\phi(Ku)$ is derived in Appendix (equations (1.4)-(1.5))
- Anti-merozoite: $p(v) = \dfrac{p_0}{1+K_v v}$ - reduction in RBC-invasion by *M*.

#### Single-strain system

**Variables:** (i) normal RBC - *x* (homeostatic maintenance, depletion by invading merozoites); (ii) Infected iRBC - *y* , free merozoites $M = r\,y$ (r-replication factor); (iii) specific immune effector (Ab) *u* ([1], [2]); binds to iRBC (prevents sequestration and clears y)

**Processes:**

1. **RBC Invasion**: newly invaded RBC $Y = \left(1 - e^{-pM/x}\right)x$, *p* – probability of invasion
2. **Immune clearing**:
   a. Innate (febrile): sigmoid $f\left(Y\right) = \dfrac{1}{1 + \left(Y / y_F\right)^p}$ - density-dependent surviving fraction of Y with febrile threshold $y_F$
   o Adaptive *u*: surviving fraction of newly invaded RBC $\phi\left(K\,u\right)Y$ , K – affinity. Clearing function $\phi\left(Ku\right)$ is derived in Appendix (equations (1.4)-(1.5))
3. **Immune stimulation**: homeostatic maintenance of baseline *u* and its stimulation (growth/proliferation) by *y* .

The system is run in discrete time steps $\Delta t = 48$ h (merozoite replication cycle), as map of current state $\left(x, y, u\right)$ at time *t* (the end of cycle) to next state $\left(x', y', u'\right)$, at $t + \Delta t$

$$x' = \underbrace{\left(1-\delta\right)x_0}_{\text{production}} + \underbrace{\delta x}_{\text{loss}} - \underbrace{Y}_{\text{merozoite invasion}}$$

$$y' = \underbrace{\phi\left(u\right) f\left(Y\right)}_{\text{immune survival}} Y + \underbrace{S}_{\text{source/inoculae}}$$

$$u' = \underbrace{\left(1-\delta_u\right)u_0}_{\text{baseline maintenance}} + \underbrace{\delta_u u}_{\text{loss}} + \underbrace{\sigma\left(y, u\right)}_{\text{stimulation}}$$

**Table 1:** Model equations (single-variant, single-strain case)

| Compartment | Equations | Auxiliary variables, triggers |
|---|---|---|
| RBC | $x' = (1-\delta)x_0 + \delta x - Y$ | RBC invasion: $Y = \left(1-e^{-pM/x}\right)x$ |
| iRBC | $y' = f(Y)\phi(u)Y + S$ - source | Free merozoites: $M = r\,y$<br>Immune clearing (iRBC survival fraction)<br>Innate: $f(Y) = \frac{1}{1+(Y/Y_T)^m}$, m – Hill exponent<br>Specific (Ab): $\phi(u)$ |
| Specific IE | $u' = (1-\delta_u)u_0 + \delta_u u + \sigma(u,y)$ | immune stimulation: $\sigma(y,u) = \frac{\sigma_0 y}{y+y_T}\cdot\frac{u}{1+u/u_T}$ |

#### Multi-strain system

Parasite types labeled by $i = 1, 2, \ldots$

- **Table 2:** Dynamic variables and functions used in multi-variant

| Variable | Notation |
|---|---|
| Uninfected red blood cells [μL$^{-1}$] | $x$ |
| Merozoites | $M_1,\ M_2,\ldots,M_n$ |
| Newly invaded RBC [μL$^{-1}$] | $Y_1,\ Y_2,\ldots,Y_n$ |
| Sequestered iRBC [μL$^{-1}$] | $y_1, y_2,\ldots,y_n$ |
| Anti-EMP (VSA) effectors | $u_1,u_2,\ldots,u_n$ |
| Anti-MSP effector | $v$ |

- **Table 3:** Model equations (single-strain case)

| Compartment | Equations | Auxiliary variables; immune triggers |
|---|---|---|
| RBC | $x' = (1-\delta)x_0 + \delta x - Y$ | $M_i = \sum_j a_{ij} r_j\, y_j$ ; total<br>$M = \sum_{i=1}^{n} M_i$ |
| Sequestered iRBC | $y_i' = f(Y)\phi(u_i)Y_i$ | $Y_i = \frac{M_i}{M}Y$ ; total $Y = \left(1-e^{-pM/x}\right)x$ |
| u- effector | $u_i' = (1-\delta_u)u_0 + \delta_u u_i + \frac{\sigma_u Y_i}{Y_i + y_T}\cdot\frac{u_i}{1+u/u_T}$ | $Y_i = \sum_m K_{im}\left[\alpha Y_m + (1-\alpha)y_m\right]$<br>$u = \sum_i u_i$ |

**Table 4:** Parameter names and values/ranges used in the model

| Parameter Definition | Notation | Value range |
|---|---|---|
| Normal red blood cell level [$\mu L^{-1}$] | $x_0$ | $5 \times 10^6$ |
| Red blood cell loss rate (survival fraction)[1] | $\delta$ | 0.98 |
| Parasite clearing cut off [$\mu L^{-1}$] | $y_C$ | $10^{-6}$ |
| Parasite replication rate [per cycle 1-2 days] | $r_i$ | 6 -12 |
| Immune stimulation threshold | $y_T$ | 0.001 - 0.003 |
| Baseline immune level (dimensionless) | $u_0$ | .05-.1 |
| | $v_0$ | .01-.05 |
| Immune efficiency (affinity) | $K_{ij}$ | 0-1 |
| Saturation threshold for immune growth | $u_T, v_T$ | 20-50 |
| Maximal proliferation rates | $\sigma_u$ | 2-4 |
| | $\sigma_v$ | 1.2-1.8 |
| Immune memory (loss rate/cycle) [2] | $\delta_u$ | 0.9 - 0.95 |
| | $\delta_v$ | .9-.99 |

#### Genetically structured parasite

Parasite makeup: collection of var-genes

| Locus | 1 (growth) | 2 (anti-erythrocyte) |
|---|---|---|
| 1 | $r_1$ | $u_1$ |
| 2 | $r_2$ | $u_2$ |
| … | … | … |

##### AV-switching

Different patterns of mutation matrix: $A = (a_{ij})$, $\sum_{i=1}^{n} a_{ij} = 1$ .

1) Uniform $a_{ij} = \varepsilon$ ($i \neq j$), with rate $\varepsilon = 10^{-9} - 10^{-4}$ $a_{ii} = 1 - \sum_{j \neq i} a_{ji}$

2) Sequential band-diagonal $A = (a_{ij})$ , $a_{ij} = \varepsilon$ ($|j-i|=1$) or 0 ($|j-i|>1$).

3) Structured mutability based on network (graph) with links $\{i \to j\}$

[1] Based on 100-day life-span of RBC
[2] Based on immune (Ab) duration of 40-100 days

4) Structured mutability based on genotypes (alleles on different loci control basic phenotypes growth, antigenic types, etc.),

So clone $(i, j, k)$ has growth - $r_i$, anti-RBC effector - $u_j$, anti-merozoite effector - $v_k$. Such structured quasi-species can be divided into cross-reacting groups.

#### Input parameters

Strain AV-diversity: n= # distinct AV types. Effective parasite replication $r = (r_1; r_2; ...)$ ; switching/mutability $\varepsilon$ . Max immune stimulation proliferation ( $\sigma_u; \sigma_v$ ), immune thresholds $y_T; M_T$ ; memory loss $\delta_u; \delta_v$, memory rates (survival fractions) $\delta_u; \delta_v$ .

#### Estimated outputs

Individual histories and "host ensembles" with various inputs, including (i) random AV-switching rate and pattern $A = A(\varepsilon)$ ; (ii) combinations of uncertain in-host parameters.

Descriptive ensemble statistics include:

1. Observed history duration
2. Max parasitemia, # peaks
3. Fast and slow clearing slopes
4. Virulence (=max RBC loss), cumulative RBC-loss/host mortality
5. Transmissibility based on parasitemia-duration patterns

#### Immune clearing and stimulation

Anti-malaria immunity involves different types of antigens and immune effectors (Abs), most important are anti-merozoite MSP-effectors ($v_1; v_2; \ldots$ ), and anti-erythrocyte EMP effectors ($u_1; u_2; \ldots$ ). Both effector types bind to respective antigens: $u_i$ - to iRBC $y_i$ (and cross-reacting $y_j$ ), while $v_i$ binds to free merozoites $M_i$ , $M_j$ . We want to estimate surviving fraction $\phi(u)$ of newly invaded iRBC $Y$, as function of immune effector *u*. The derivation employs continuous Ab-binding process (DE). Two possible approaches are (i) binding-dissociation kinetics, and transition 'free' to 'opsonized' iRBC, $y \rightleftharpoons z$ ; (ii) dynamics of opsonization fraction of iRBC, $0 < s(t) < 1$, and the resulting survival probability $\phi(s)$

**Binding- dissociation kinetics.** We use standard reaction kinetic for free and bound iRBC populations $y(t)$ $z(t)$

$$y \underset{k_-}{\overset{k\,u}{\rightleftharpoons}} z \overset{\nu}{\rightarrow} \qquad (0.1)$$

with binding rate $k\,u$, proportional to concentration $u(t)$, and fixed dissociation $k_-$. Opsonized population *z* is cleared (spleen) at a rate $\nu$, and is removed by the end of the cycle (no merozoite release). So only free (unbound) *y* are left to release merozoites for the next cycle.

We want to estimate the surviving fraction of initial population $Y$ by the end of the cycle $0 < t < T$ (= 48 hrs). The DE system for reaction kinetics (1.1),

$$\begin{aligned} \frac{dy}{dt} &= -k\,u\,y + k_- z; \qquad y(0) = Y \\ \frac{dz}{dt} &= k\,u\,y - (k_- + \nu) z; \quad z(0) = 0 \end{aligned} \qquad (0.2)$$

initialized at values $y(0) = Y = \left(1 - e^{-pm/x}\right)x$, $z(0) = 0$, all newly invaded iRBC unbound by *u*. After *rescaling* over dissociation rate $k_-$ , $k \rightarrow K = k / k_-$ - affinity, $\nu \rightarrow \nu / k_-$, assuming near constant (mean) effector level *u* over the cycle, we write solution of (1.2) via eigenvalues of matrix

$$A = \begin{bmatrix} -Ku & 1 \\ Ku & -(1+\nu) \end{bmatrix}$$

$$\lambda_{1,2}(K\,u,\nu) = -\frac{1+K\,u+\nu}{2} \pm \sqrt{\left(\frac{1+K\,u+\nu}{2}\right)^2 - \nu\,K\,u} \quad (0.3)$$

The survival fraction is given by

$$\frac{y(t)}{Y} = \phi(Ku) = \frac{(K\,u+\lambda_2)e^{\lambda_1} - (K\,u+\lambda_1)e^{\lambda_2}}{\lambda_2 - \lambda_1} \quad (0.4)$$

Function $\phi(Ku)$ predicts (approximately) surviving fraction of iRBC by the end of cycle in terms of dimensionless $K\,u$, and relative clearing rate $\nu = \nu / k_-$.

For multi-strain systems with immune effectors $\{u_1;...;u_n\}$ and cross-reactive matrix $K = (K_{ij})$ we use approximate clearing function (1.3)-(1.4) applied to each strain $y_i$, so

$$\frac{y_i}{Y_i} = \phi\left(\sum\nolimits_{j=1}^{n} K_{ij}u_j;\nu;T\right) = \phi\left(\sum\nolimits_{j=1}^{n} K_{ij}u_j\right) \quad (0.5)$$

**Opsonisation process:** the dynamic variable here is mean coverage level $s(t)$ of iRBC bound by Ab *u*. The binding-dissociation kinetics gives DE

$$\frac{ds}{dt} = k\,u(1-s) - k\_s \quad (0.6)$$

Its equilibrium level $s^* = \frac{Ku}{1+Ku}$ , determines mean survival probability as sigmoid function

$\phi(s) = \frac{1-s^m}{1+s^m}$ (Figure 1).

The resulting survival probabilities in both systems, binding-dissociation (i) and opsonisation (ii) are shown in Figure 2. They are qualitatively similar for both systems.

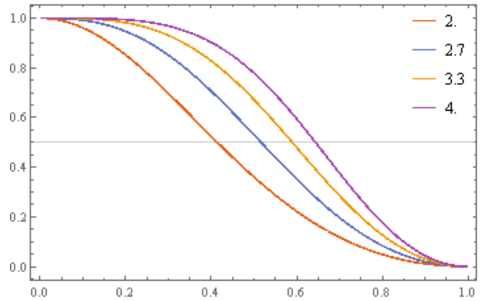

Figure 1: Survival functions $\phi(s)$ for seral values $2 \le m \le 4$

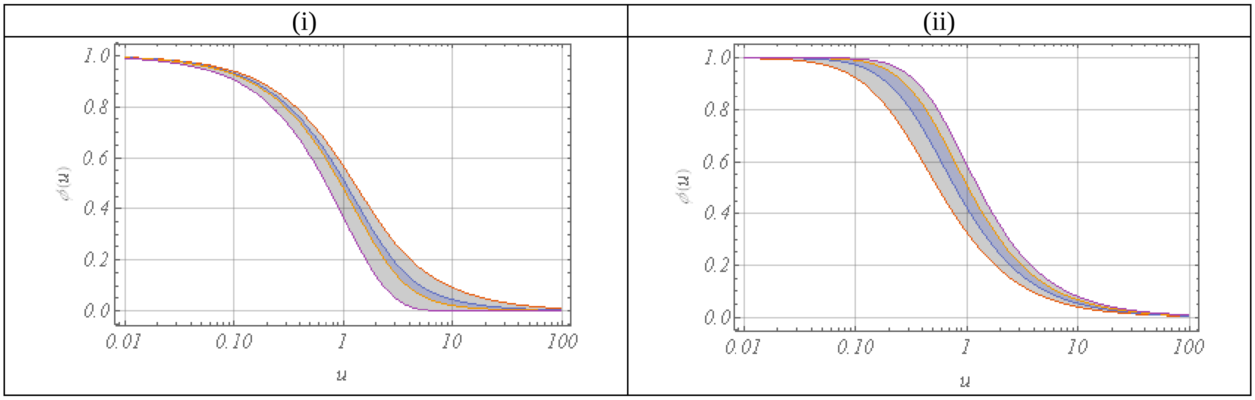

Figure 2: Survival fraction $\phi(u)$ in 2 cases: system (i) for clearing range $0 < \nu < \infty$ (top-to-bottom) – shaded; system (ii) for range $2 \le m \le 4$

**Anti-merozoite** Abs $v = (v_i)$ have short time to invade RBC, so the clearing is less important ($\nu = 0$), but only unbound fraction *m* of freshly released merozoite $M = r\,y$ can invaded RBC efficiently. We estimate this fraction from equilibrium M- kinetics

$$\frac{dm}{dt} = -k\,v\,m + k_{\_}(M - m)$$

Hence, $m^* = \frac{M}{1 + K\,v}$ , with affinity $K = K_v = \frac{k}{k_{\_}}$ , or $\frac{M_i}{1 + \sum_j K_{ij} v_j}$ - for cross-reacting suite $(v_j)$ .

**Immune stimulation**/growth for $u_i$ (or $v_i$) is determined by antigenic load $Y_i$ ($M_i$), or cross-reacting $\sum_j K_{ij} Y_j$ .

Variable *Y* stimulates proliferation of immune B-cells, their conversion to secreting (plasma) cells and release of antibodies (effector *u*). We lump all these processes into a single immune stimulation function

$$\sigma(Y) = \frac{\sigma_M Y}{Y + y_T}$$

with maximal proliferation rate $\sigma_M$ , and sensitivity threshold $y_T$ . So $\sigma(Y)$ changes from a 'decay mode' ( $\sigma(Y) < 1$ at low Y) to the 'growth mode' ( $\sigma(Y) > 1$), as $Y$ passes a critical threshold.

The exponential growth of *u* driven by factor $\sigma(Y) > 1$ cannot proceed indefinitely, but it should slow down when B-cell populations and effector *u* reach sufficiently high levels. While multiple factors and mechanisms control proliferation/growth of immune effectors, in our scheme we implement it through a simple density threshold effect, so it exponentially for low $u < u_T$ (threshold), and saturates at high levels by density factor $\frac{u}{1 + u / u_T}$. The combined immune stimulation function takes the form

$$\sigma(Y,u) = \frac{\sigma_M Y}{Y + y_T} \cdot \frac{u}{1 + u / u_T} \tag{7}$$

We also allow for slow waning of immune effector *u* at a certain rate $0 < \delta_u < 1$, due to Ab degradation and memory loss.

1. Struik SS, Riley EM. Does malaria suffer from lack of memory? *Immunol Rev.* 2004;201:268-290.
2. Eckhoff P. P. falciparum infection durations and infectiousness are shaped by antigenic variation and innate and adaptive host immunity in a mathematical model. *PloS one.* 2012;7(9):e44950.
